# Collagen receptor GPVI-mediated platelet activation and pro-coagulant activity aggravates inflammation and aortic wall remodelling in abdominal aortic aneurysm

**DOI:** 10.1101/2023.11.20.567851

**Authors:** T Feige, A Bosbach, KJ Krott, J Mulorz, M Chatterjee, J Ortscheid, E Krüger, I Krüger, W Ibing, M Grandoch, MU Wagenhäuser, H Schelzig, M Elvers

**Author notes:** contributed equally (T Feige, A Bosbach, KJ Krott). corresponding author **E-Mail of Corresponding author:** Margitta Elvers, Ph.D.

## Abstract

Platelets play an important role in cardio- and cerebrovascular diseases. Abdominal aortic aneurysm (AAA) is a highly lethal, atherosclerotic-related disease with characteristic features of progressive dilatation of the abdominal aorta and degradation of the vessel wall accompanied by chronic inflammation. Platelet activation and pro-coagulant activity play a decisive role in the AAA pathology as they might trigger AAA development in both mice and men. The present study investigated the impact of the major platelet collagen receptor glycoprotein (GP)VI in cellular processes underlying AAA initiation and progression. Genetic deletion of GPVI offered protection of mice against aortic diameter expansion in experimental AAA. Mechanistically, GPVI deficiency resulted in decreased inflammation with reduced infiltration of neutrophils and platelets into the aortic wall. Further, remodelling of the aortic wall was improved in absence of GPVI, indicated by reduced MMP2/9 and OPN plasma levels and an enhanced α-SMA content within the aortic wall, accompanied by reduced cell apoptosis. As a result, an elevation in intima/media thickness and elastin content were observed in GPVI-deficient PPE mice, coursing a significantly reduced aortic diameter expansion and reduced aneurysm incidence. In AAA patients, enhanced plasma levels of soluble GPVI and fibrin, besides fibrin accumulation within the intraluminal thrombus (ILT) suggested that GPVI might serve as a biomarker and mediator in fibrin-supported stabilization of the ILT. In conclusion, our results emphasize the potential need for a GPVI-targeted anti-platelet therapy to reduce AAA initiation and progression, as well as to protect AAA patients from aortic rupture.

**Translational perspective:** Abdominal aortic aneurysm (AAA) is an atherosclerotic-related, cardiovascular disease (CVD) with high mortality. The impact of platelets in different cellular processes underlying AAA initiation and progression remains unclear.Therefore, we analysed the role of the major platelet collagen receptor GPVI in the pathogenesis of AAA. Results from platelet depleted mice and patients with AAA revealed a significant contribution of GPVI to the inflammatory response and remodelling process of the aorta. Further, elevated accumulation of fibrin, a recently identified ligand of GPVI in the intraluminal thrombus (ILT) and in the plasma of AAA patients, suggests that GPVI binding to fibrin plays a role in ILT formation and probably stabilization of the abdominal aorta. Furthermore, increased levels of sGPVI suggest that GPVI might serve as a clinical biomarker for AAA. Thus, therapeutic targeting of GPVI-mediated platelet activation might be an effective anti-thrombotic strategy for AAA patients.

## 1. Introduction

Atherosclerotic diseases are a leading cause of cardiovascular related deaths worldwide (1). Abdominal aortic aneurysm (AAA) is an atherosclerotic-related disease with a high mortality rate if untreated. The prevalence of AAA, defined as an aortic diameter of ≥ 3 cm is age-dependend. Further, well-known risk factors are male gender, family history of AAA, smoking, hypertension, hypercholesterolemia, obesity and cardiovascular diseases. The global prevalence of AAA is ∼ 0.92 % among persons aged 30-79 translating to at total of 35.12 million AAA cases worldwide in 2019 (2). With enhanced AAA diameter, the risk of fatal rupture inrceases exponentially leading to high mortality in AAA. Despite great efforts have been made to understand the underlying pathological mechanisms of AAA, the only treatment option to date is endovascular and open surgical repair, once the aortic diameter exceeds defined cut-off values. Although there are some promising candidates for non-invasive drug-based therapies, clinical implementation of such therapies are lacking besides, prospective randomized trials have either not demonstrated expected therapeutic efficacy or have not been performed to date (3–7). The characteristic progressive dilatation of the abdominal aorta is accompanied by degradation and remodelling, weakening of the vessel wall due to chronic inflammation. This is coursed by leukocyte infiltration and the extracellular matrix degradation by the activation of proteolytic enzymes (5, 6, 8, 9).

Platelets, besides their primary role in thrombosis, haemostasis, and maintaining vascular integrity, are prompt drivers of inflammation as they release various cytokines, chemokines, growth factors and biological modulators upon activation. Thereby, platelets contribute to inflammation, angiogenesis, cell survival as well as vascular and tissue repair/regeneration (10). In contrast to various cardiovascular diseases, relatively little is known about the impact of platelets in the pathological development of AAA, although experimental and clinical data strongly suggest that platelet activation may play a pivotal role in AAA initiation and progression.

Support for an active contribution of platelet response in the development and progression of AAA in humans comes from different clinical trials. Anti-platelet medication of small-sized AAA in which an intraluminal thrombus (ILT) is seldom present have not shown any significant difference. Nevertheless, other clinical trials with AAA sized above 35 and 39 mm have provided evidence of a beneficial effect of anti-platelet medication with low-dose aspirin, since aneurysm size was decreased and less frequent need for surgical repair was reported (11). Similary, in various preclinical animal models such as the angiotensin-II (Ang-II) and the pancreatic porcine elastase (PPE) infusion model, anti-platelet treatment or genetic ablation of specific platelet proteins have demonstrated a beneficial impact on AAA rupture rates, AAA growth and inflammation (12–18).

Recently, our group could demonstrate a pivotal role of platelet activation in the experimental murine PPE model suggesting a crosstalk of platelets with target cells such as macrophages and fibroblasts leading to inflammation, stiffening of the aortic wall and AAA growth (19). In this study, we observed a major contribution of platelets in the inflammatory response and in the remodelling process of the aortic wall in experimental AAA, resulting from enhanced platelet activation and pro-coagulant activity. Elevated platelet reactivity and pro-coagulant response was also documented in AAA patients, strongly supporting the potential relevance of anti-platelet therapy to reduce AAA progression and rupture in AAA patients (19). In this study we reported for the first time a pathophysiological relevance of the platelet-specific glycoprotein (GP)VI, since GPVI activation of platelets induced enhanced gene expression of acute phase cytokines and MMP9 in macrophages and IL-6 release from aortic fibroblasts. However, elevated platelet activation was due to an increase in GPVI activation at day 28 after elastase infusion into mice (19). Nevertheless, GPVI has been shown to represent a key receptor in acute coronary syndrome (ACS) and is a useful biomarker for the early detection of atherosclerotic diseases such as ACS and ischemic stroke. Further, platelet GPVI may contribute to risk stratification and prediction of clinical outcome in patients with atherosclerotic diseases. In addition, therapeutical implications of targeting GPVI in mice and men have been described such as Revacept, a GPVI–Fc fusion protein that binds to collagen preventing binding of platelet GPVI to collagen or by using GPVI-blocking antibodies (20–25). Although limited data, evaluating the role of GPVI in the context of AAA are available, there is still a significant lack of understanding on how platelets participate in AAA development and progression and whether, when and how GPVI contributes to these processes. However, more mechanistic insights into the specific role of platelets and GPVI in AAA pathology are urgently needed to allow the development of appropriate antiplatelet targets. We already showed recently that platelets strongly affect cellular processes in AAA that foster AAA progression (19). Here, we now provide strong evidence that GPVI plays a dominant role in these processes. Thus, there is a strong need to evaluate GPVI as a potential target for novel drug-based AAA thaperies; in particular, since there is no drug-based therapy to treat AAA patients available so far according to European guidelines.

## 2. Methods

### 2.1 Human AAA samples and study approval

Fresh citrate-anticoagulated blood (BD– Vacutainer^®^; Becton, Dickinson and Company; #367714) was collected from abdominal aortic aneurysm (AAA) patients. Blood samples of healthy volunteers with an average age of 61.7 ± 1.57 years served as controls. Control blood samples were obtained from the blood donor center at the University Hospital of Duesseldorf, Germany. Human intraluminal thrombus (ILT) tissue of AAA patients and tissue of arterial thrombi (AT) from thrombectomy patients were obtained from patients, which underwent surgery in the department of Vascular- and Endovascular Surgery at the University Hospital Duesseldorf. All participants provided written informed consent to participate in this study according to the guidelines of thr Ethics Committee of the University Clinic of Duesseldorf, Germany (2018-140-kFogU, study number: 2018064710; biobank study number: 2018-222_1; MELENA study 2018-248-FmB, study number: 2018114854; 4669R, study number: 2014042327). This study was conducted according to the Declaration of Helsinki and the International Council for Harmonization Guidelines on Good Clinical Practice. All procedures were approved by the Ethics Committee of the University Hospital of Duesseldorf, Germany.

### 2.2 Human plasma preparation

For plasma preparation, citrate-anticoagulated blood human blood samples were centrifuged at 1,000 x *g* for 10 min at RT. After centrifugation, the platelet-free plasma (PFP) was collected and stored at -70 °C until use.

### 2.3 Animals

Pathogen-free mice with GPVI (*Gp6^-/-^*) ablation were provided by J. Ware (University of Arkansas for Medical Sciences, Little Rock, AR, USA) and backcrossed to C57BL/6J mice. All experiments were conducted with male mice aged 10–12 weeks. The mice were maintained in an environmentally controlled room at 22 ± 1 °C and a 12 h day-night cycle. Mice were housed in Macrolon cages type III with *ad libitum* access to food (standard chow diet) and water. All animal experiments were conducted according to the Declaration of Helsinki and were approved by the Ethics Committee of the State Ministry of Agriculture, Nutrition and Forestry State of North Rhine-Westphalia, Germany (permit number: 81-02.05.40.21.041 and 81-02.4.2018.A409).

### 2.4 Experimental AAA mouse model

The porcine pancreatic elastase (PPE) perfusion model is an experimental mouse model to study AAA development and progression *in vivo* by infusion of porcine pancreatic elastase into the abdominal infrarenal aorta. To induce AAA formation, mice were anesthetized with 2–3 % isoflurane and received an additional subcutaneous (s.c.) injection of buprenorphine (0.1 mg/kg body weight) (Temgesic, Eumedica) 30 min prior PPE-surgery. Thereby, anaesthesia was continuously maintained andmonitored throughout the procedure. The PPE surgery was conducted as prevoisly described (15, 19).

In short, the infrarenal aortic segment was infused with elastase at 2.5 U/mL for 5 min. After surgery all mice received an additional subcutaneous (s.c.) injection of buprenorphine (0.1 mg/kg body weight) every 6 h during the daylight phase and during the night phase via drinking water (0.3 µg/mL) for 3 days for pain relief. For aortic tissue collection mice were euthanized via cervical dislocation under deep isoflurane anaesthesia at day 7, 14 and 28 post PPE surgery.

### 2.5 Ultrasound imaging

AAA development and progression in PPE-operated mice was monitored via ultrasound measurements. The maximum dilation of the abdominal aorta within the aneurysm segment was measured prior to (baseline) and at day 7, 14, 21 and 28 after PPE surgery. For ultrasound imaging mice were anesthetized with 2–3 % isoflurane and placed onto a heating plate at 37 °C. All ultrasound measurements were performed using a Vevo 3100^®^ High-Resolution In Vivo Micro-Imaging System (VisualSonics). For monitoring of the inner aortic diameter progression a standardized imaging algorithm with longitudinal B– Mode images of the systolic phase was used.

### 2.6 Preparation of murine aortic tissue

Mice were euthanized via cervical dislocation under deep isoflurane anaesthesia at day 7, 14 and 28 post PPE surgery. After euthanasia, the thorax was opened and the heart was punctured with a butterfly cannula at the apex cordis of the left ventricle. The vascular system was subsequently perfused under constant pressure with approximately 20 mL cold heparin solution (20 U/mL, 4 °C, #2047217, Braun) to avoid clot formation. Afterwards the abdominal aorta was removed and fixed in 4 % paraformaldehyde (PFA, #P087.5, Carl Roth) for 24 h at 4 °C. After dehydration in an ethanol gradient, the aortic tissue was incubated for at least 12 h in Roti^®^Histol (#6640.4, Carl Roth) at RT and embedded in paraffin (#P3558, Sigma-Aldrich).

### 2.7 Murine blood collection and plasma preparation

Murine whole blood was collected in 300 µL heparin solution (20 U/mL). Blood cell counts, including platelet count and mean platelet volume (MPV), were analysed using a hematology analyzer (Sysmex KX21N, Norderstedt). For plasma preparation blood was collected into EDTA coated Microvettes^®^ (#20.1341, Sarstedt). Samples were centrifuged at 2,500 x *g* for 5 min at 4 °C. Plasma samples were stored at -70 °C until use.

### 2.8 Flow cytometry

For flow cytometric analysis of murine platelets, heparinized murine whole blood was washed 3 times with Tyrode’s buffer (137 mM NaCl, 2.8 mM KCl, 12 mM NaHCO_3_, 0.4 mM NaH_2_PO_4_ and 5.5 mM glucose, pH 6.5) by centrifugation at 650 x *g* for 5 min. After final washing the samples were resuspended in murine Tyrode’s buffer supplemented with 1 mM CaCl_2_. For measuring platelet activation a two-color analysis was performed using specific antibodies for P-selectin (CD62P, Wug.E9-FITC, #D200, Emfret Analytics) and active integrin α_IIb_β_3_ (JON/A-PE, #D200, Emfret Analytics) in a final ratio of 1:10. Samples were incubated with the indicated agonists (adenosine diphosphate (ADP; #A2754, Sigma-Aldrich), collagen-related peptide (CRP; Richard Farndale, University of Cambridge, Cambridge, UK), the thromboxane A2 analogue U46619 (U46; #1932, Tocris) or PAR4 peptide (PAR4; #3494, Tocris)) and the respective antibodies for 15 min at 37 °C in the dark. Reaction was stopped by adding 300 µL of PBS to all samples. For analyzing glycoprotein (GP) exposure, washed whole blood samples were incubated with specific antibodies against integrin α_5_ (CD49e, Tap.A12-FITC, # M080-1, Emfret Analytics), GPIbα (CD42b, Xia.G5-PE, #M040-2, Emfret Analytics) or GPVI (JQ1-FITC, #M011-1, Emfret Analytics) in a ratio of 1:10 for 15 min at RT. To determine upregulation in glycoprotein surface expression, washed blood was specifically labelled for integrin β_3_ (CD61, Luc.H11-FITC, #M031-1, Emfret Analytics) at a ratio of 1:10 and stimulated with the indicated agonists. To measure phosphatidylserine (PS)-exposure on the platelet surface washed murine whole blood was resuspended in binding buffer (10 µM HEPES, 140 µM NaCl, 2.5 mM CaCl_2_, pH 7.4). All samples were labelled with a PS detecting antibody (Cy^TM^ 5 Annexin V, #559934, BD Bioseciences) at a ratio of 1:10. Samples were incubated with the indicated agonists and the antibodies for 15 min at 37 °C in the dark. Reaction was stopped by adding 300 µL of binding buffer (0.1 M HEPES, 1.4 M NaCl, 25 mM CaCl_2_, pH 7.4). All samples were labelled for 15 min at RT. To analyze platelet-leukocyte aggregate formation, washed murine whole blood was specifically labelled for the leukocytes (CD45, 30-F11-APC, #559864, BD Biosciences), neutrophils (Ly6G, 1A8-APC, #127614, BD Biosciences), T cells (CD3, 17A2-APC, #100235, BioLegend), macrophages or granulocytes (CD14, Sa14-2-APC, #123311, BioLegend and CD11b, M1/70-APC, #553312, BD Biosciences) at a ratio of 1:10. GPIbα (CD42b, Xia.G5-PE, #M040-2, Emfret Analytics) served as a platelet specific marker. All samples were labelled for 15 min at RT in the dark. For analyzing human platelets, whole blood samples were diluted at a ratio of 1:10 in human Tyrode’s buffer (137 mM NaCl, 2.8 mM KCl, 12 mM NaHCO_3_, 0.4 mM NaH_2_PO_4_ and 5.5 mM glucose, pH 6.5) and specifically stained for 15 min at RT using an antibody against GPVI (HY101-PE, #565241, BD Pharmingen^TM^) at a ratio of 1:10. All samples were analysed using a BD FACSCalibur™ flow cytometer.

### 2.9 Flow cytometric based multiplex analysis

Plasma level of the circulating inflammatory cytokines IL1α, IL1β, IL6, IL10, IL17A, INF, and GM-CSF in PPE-operated mice were determined using a flow cytometry based cytometic bead multiplex analysis (LEGENDplex^TM^ Mouse Inflammation panel (13-plex), #740150, BioLegend). All measurements were conducted according to the manufacturer’s instructions by using a BD FACSLyric^TM^ flow cytometer (26).

### 2.10 Enzyme-Linked Immunosorbent Assay (ELISA)

For analysis of murine plasma samples an OPN (#DY441, R&D Systems), a neutrophil elastase (ELA2) (#DY4517-05, R&D Systems), MMP2 (#MMP200, R&D Systems) and MMP9 (#MMPT90, R&D Systems) ELISA were performed. For the analysis of AAA patient plasma samples a human fibrin (#MBS265263, MyBioSource) and a human soluble GPVI (#MBS9390142, MyBioSource) ELISA were used. All ELISA were conducted according to the manufacturer’s instructions.

### 2.11 Immunofluorescence staining of murine aortic tissue

Paraffin-embedded aortic tissue from PPE-operated mice was sliced into 5 µm sections using an automatic microtome (Microm HM355, Thermo Fisher Scientific). Prior to staining, all sections were deparaffinized and hydrated. For antigen unmasking the tissue sections were heated at 300 W in citrate buffer (pH 6.0) for 10 min. For staining of macrophages and platelets the aortic tissue sections were blocked for 1 h at RT with protein blocking solution (#X0909, Dako). After blocking, the sections were specifically stained for macrophages (Mac3 (CD107b), #550292, BD Pharmingen^TM^, 1:50) or platelets (GPIbα (CD42b), #M042-0, Emfret Analytics, 1:50) at 4 °C overnight. Respective IgG primary antibody served as negative controls (#C301, Emfret Analytics, 1:50). The sections were incubated with a biotinylated secondary antibody (#BA-9400, Vector, 1:200) for 1 h at RT and afterwards with a streptavidin eFluorTM 660 conjugate (#50-4317-80, Thermo Fisher Scientific, 1:20) for 30 min at RT. For staining of neutrophils (Ly6G, #551459; BD Pharming^TM^, 1:100), osteopontin (OPN, #ab1841440, abcam, 1:50), smooth muscle actin (αSMA, #ab5694, abcam, 1:50) and cleaved caspase3 (Casp3, #9661, Cell Signaling, 1:50) respectively, the aortic tissue sections were blocked with DPBS (containing 0.3 % Triton x-100 and 5 % goat serum) for 1h at RT. After overnight incubation with the respective primary antibodies at 4 °C, the section were stained with a highly crossed-absorbed Alexa Fluor^TM^ Plus 555 conjugated secondary antibody (#A32794, Invitrogen^TM^, 1:200) for 1 h at RT. Respective IgG antibodies served as negative control (#3900S, Cell Signaling, 1:5). For visualizing nuclei, all tissue sections were stained with DAPI (4′, 6-Diamidine-2′-phenylindole dihydrochloride, #10236276001, Roche, 1:3000). For image generation an Axio Observer. D1 microscope (Zeiss) was used. Tissue section of PPE-operated were analysed as percentage content normalised to the total aortic tissue, using Image J (version 1.53t) by applying the preinstalled “*Default*” threshold. Per cross-section 4 representative fields were analysed. For quantification of the aortic elastin content the mean fluorescence intensity (MFI) was determined using the ZEN software (Zeiss, blue edition, version 3.3.). The MFI of each section was normalised to the total area of the aortic tissue.

### 2.12 Histology of murine aortic tissue

Paraffin-embedded aortic tissue from PPE mice was sliced into 5 µm sections using an automatic microtome (Microm HM355, Thermo Fisher Scientific). Hematoxylin/eosin (H/E) staining of murine aortic tissue was conducted as described elsewhere (27). The intima/media thickness of PPE-operated mice was analysed by measuring the aortic media thickness at 10 distinct positions within the aneurysm segment using the ZEN software (Zeiss, blue edition, version 3.3.). The modified Verhoeff van Gieson (VVG) staining in murine aortic tissue was performed according to the manufacturers instructions (Elastic stain, #HT25A-1KT, Sigma-Aldrich). For analyzing elastin fragmentation the number of elastin breaks was determined using the ZEN software (Zeiss, blue edition, version 3.3.). Moreover, picrosirius red staining of murine aortic tissue was conducted as described elsewhere (28). To determine the collagen composition within the aneurysm segment the collagen density was anaylzed under polarized light using an Axio Imager 2 (Zeiss). The collagen content was determined as percent normalised to the total aortic tissue, using the Image J (version 1.53t) software by applying the preinstalled “*Default*” threshold.

### 2.13 Histology of human thrombus tissue

For Martius Scarlet Blue (MSB) staining of human thrombus tissue, paraffin-embedded sections were hydrated and incubated in Bouin’s Solution (#HT10132, Sigma-Aldrich) for 1 h at 56 °C. All sections were washed for 10 min in tap water and for 5 min in 96 % EtOH. Afterwards, the tissue sections were stained with Napthol yellow S (#sc-215544, Santa Cruz Biotechnology) for 5 min at RT. After staining, the sections were washed in dH_2_O for 5 min and subsequently stained using a Crystal scarlet solution for 10 min at RT. For differentiation, all sections were incubated in 1 % phosphotungstic acid (#100583, Merck Millipore) for 10 min at RT, followed by an incubation with Methyl blue (#M5528, Sigma-Aldrich) for 15 min at RT. Tissue section were washed once in dH_2_O and for 10 sec. in 1 % acetic acid. Afterwards, the tissue sections were dehydrated and embedded using a Roti^®^-Histokitt mounting medium. For immunohistochemical staining of platelets in human thrombus tissue, paraffin-embedded sections were deparaffinized and hydrated. For antigen unmasking the tissue sections were heated at 300 W in citrate buffer (pH 6.0) for 10 min. The sections were washed 3 times in Tris-buffered saline supplemented with 0.1 % Tween^®^ 20 (TBS-T) for 5 min at RT. Tissue sections were blocked for 1 h at RT with protein blocking solution (#X0909, Dako). After blocking, the sections were specifically stained for platelets (GPIbα (CD42b), #MA5-11642, Invitrogen, 1:100) at 4 °C overnight. Respective IgG primary antibody served as control (#5415S, Cell Signaling Technology, 1:1250). For endogenous peroxidase activity blocking the sections were incubated with 0.3 % hydrogen peroxide for 15 min at RT. Afterwards, the sections were incubated with a biotinylated secondary antibody (#BA-9200, Vector Laboratories, 1:300) for 30 min at RT, followed by an incubation with an avidin-biotinylated HRP enzyme complex (#PK-6100, Vector Laboratories) for 30 min at RT. For antigen detection the tissue sections were incubated with a VIP peroxidase substrate (#SK-4600, PK-6100, Vector Laboratories) for 2 min at RT. Nuclei were visualized by incubation with Methyl green (#H-3402, Vector Laboratories) for 10 min at 60 °C. After washing in dH_2_O the sections were dehydrated and mounted using Roti^®^-Histokitt mounting medium. Tissue section of human thrombus samples were analysed as percentage content normalised to the total thrombus tissue, using Image J (version 1.53t) software by applying the preinstalled “*Default*” threshold. Per thrombus sample at least 4 representative fields were analysed.

### 2.14 Thrombus formation under flow *ex vivo* (flow chamber experiments)

Rectangular glass coverslips (24 x 60 mm) were coated with 200 µg/mL collagen (Horm^®^ collagenreagent, #1130630, Takeda Austria GmbH) at 4 °C for overnight. After coating the coverslips were blocked with 1 % BSA (in PBS) for 1 h at RT. Fresh citrate-anticoagulated human blood was pre-labeled with Mepacrine (Sigma-Aldrich) for 10 min at RT and perfused over the pre-coated coverslips using a flow chamber system (50 µm x 5 mm) under a shear rate of 1,700 s^-1^ (12.75 mL/h). Five representative images were taken using an Axio Observer. D1 microscope (Zeiss). The thrombus formation was analysed as percentage of surface coverage by using Image J (version 1.53t).

### 2.15 Statistics

Data are presented as arithmetic means ± SEM (Standard error of mean), statistical analyses were performed using GraphPad Prism 8 (version 8.4.3). The indicated sample size for each experiment reflects the number of independent biological replicates. Statistical differences were determined using one-way ANOVA with Tukey’s multiple comparisons post-hoc test, or two-way ANOVA with a Sidak’s multiple comparison post-hoc test, respectively, unpaired multiple t-test or unpaired student’s t-test. Significant differences are indicated by asterisks (*** p < 0.001; ** p < 0.01; * p < 0.05).

## 3. Results

### 3.1 GPVI-deficient mice are protected against experimental AAA progression

We induced experimental murine AAA in GPVI-deficient and control mice utilizing the porcine pancreatic elastase (PPE) model to investigate wether the genetic loss of GPVI impacted on different cellular processes contributing to the initiation and progression of AAA (**Figure 1A**). While the overall survival rate was unaltered between the groups (**Figure 1B**), we detected significantly attenuated aortic diameter progression of AAA over a 28-day time course by ultrasound tracking (**Figure 1C–D, Supplemental Figure S1A**). Reduced aortic diameter in GPVI-deficient mice was confirmed by H/E-staining at the end of the observation period of 28 days (**Figure 1E–F).** Furthermore, AAA incidence was reduced with only 36.4% of GPVI-deficient mice displaying an AAA, based on a cut-off value of 150 % from baseline diameter. In contrast, 73.3% of the wildtype PPE controls showed AAA development **(Figure 1G).** In contrast, no differences were observed with regard to body weight, platelet counts and mean platelet volume (MPV) at baseline and at different time points after elastase infusion **(Supplemental Figure 1B–D).**

**Figure 1.**
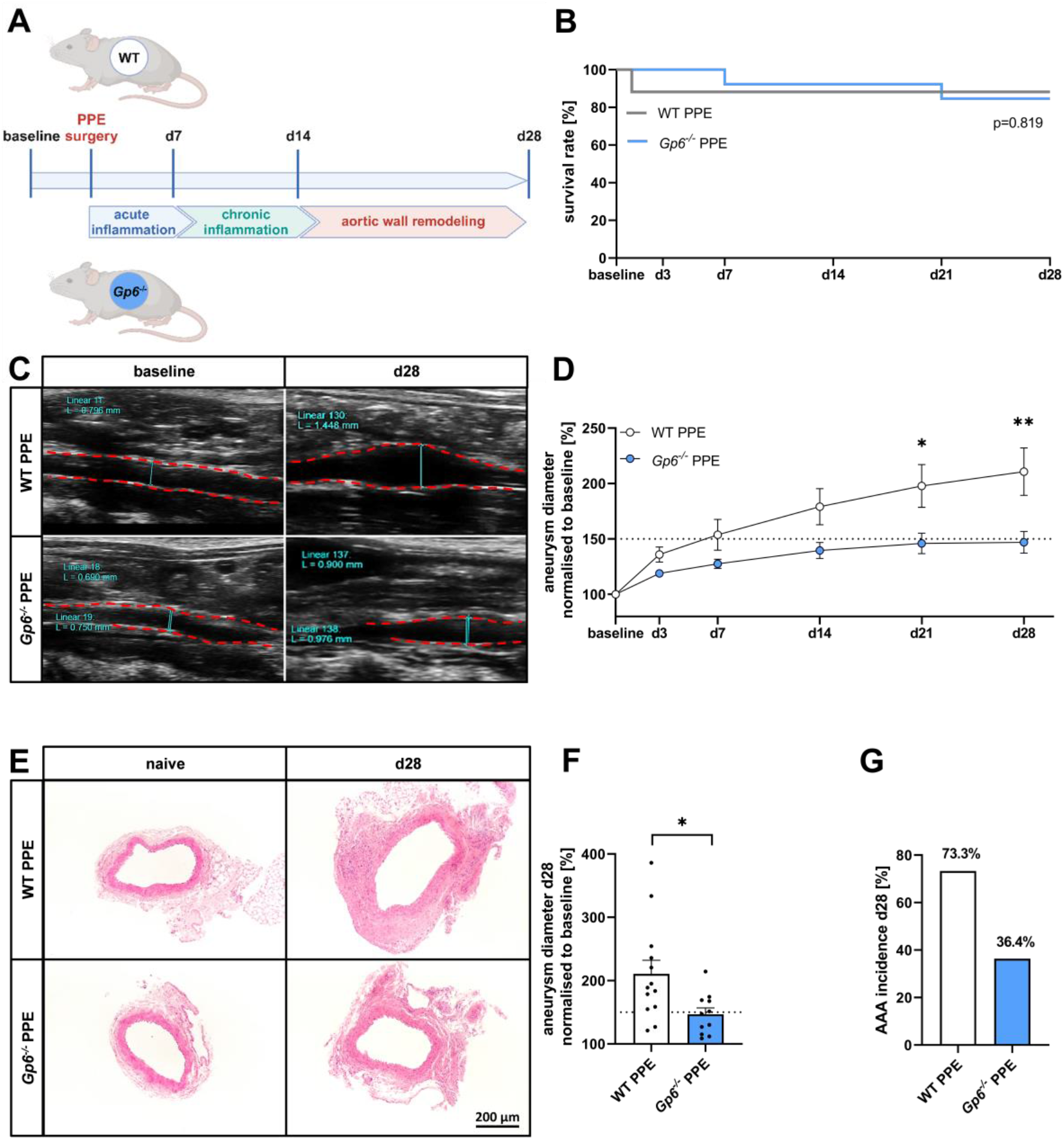
GPVI-deficient mice are protected against experimental AAA progression. **(A)** Schematic overview of experimental procedure: C57BL/6J wild type (WT PPE) and *Gp6* deficient (*Gp6^-/-^* PPE) mice underwent PPE surgery. Operated mice were sacrificed and analysed at day 7, 14 and 28 after surgery**. (B)** Survival rate of PPE-operated WT (n=17) and *Gp6^-/-^* (n=16) mice over a time period of 28 days. **(C)** Representative ultrasound images of WT (n=13) and *Gp6^-/-^* (n=11) PPE mice prior (baseline), respectively 28 days after surgery. **(D)** Aortic diameter progression of WT (n=13) and *Gp6^-/-^* (n=11) PPE mice over a time period of 28 days, analysed via ultrasound measurements. Data were normalised to baseline. **(E)** Representative overview images of the aortic tissue from WT (n=6) and *Gp6^-/-^* (n=5) PPE mice at day 28 using H/E staining. Aortic tissue from naive WT (n=6) and *Gp6^-/-^* (n=5) mice served a control. Scale bar: 200 µm. **(F)** Aneurysm diameter of WT (n=13) and *Gp6^-/-^* (n=11) mice at day 28 post PPE surgery. Data were normalised to baseline. **(G)** AAA incidence at day 28 of WT (n=13) and *Gp6^-/-^* (n=11) PPE mice revealing an aortic diameter of ≥ 150 % following 28 day after surgery. Data are represented as mean values ± SEM. Statistical analysis was performed using (**B**) a Log-rank (Mantel-Cox) test, (**D**) two-way ANOVA with a Sidak’s multiple comparisons post-hoc test and (**G**) an unpaired student’s t-test; *p < 0.05, **p < 0.01. AAA = Abdominal aortic aneurysm; ECM = Extracellular matrix; H/E = Hematoxylin-eosin; PPE = Porcine pancreatic elastase (infusion).

### 3.2 Important role of platelet GPVI in mediating platelet activation and pro-coagulant activity in AAA

Since platelet activation and pro-coagulant activity of platelets are crucial for AAA progression (19), we analysed platelet activation in GPVI-deficient and WT mice at different time points following elastase infusion (**Figure 2**). While no alterations of platelet activation and pro-coagulant activity were observed in naive (unoperated) mice (apart from GPVI defective platelet activation, **Supplemental Figure 2A and B**), we detected reduced P-selectin exposure and reduced integrin α_IIb_β_3_ activation only in response to CRP, while activation following stimulation of platelets with agonists that activate G-protein coupled receptors were unaltered at day 7 after elastase infusion (**Figure 2A).** In contrast, at day 14 and 28, reduced P-selectin exposure of GPVI-deficient platelets was not only limited to stimulation with CRP but also evident, when platelets were stimulated with ADP and the thromboxane analogue U46619. In addition, reduced P-selectin exposure of GPVI-deficient platelets was also observed following stimulation of the thrombin receptor PAR4 at day 28. In contrast, integrin α_IIb_β_3_ activation was not altered following G-protein coupled receptor stimulation at day 28 but reduced at day 14 following stimulation of platelets with ADP alone (**Figure 2B–C**). Basal pro-coagulant activity of non-stimulated GPVI-deficient platelets-as shown by Annexin-V binding-was reduced at day 7 but not at later time points (**Figure 2A–C**). Furthermore, platelet activation with CRP and PAR4 peptide resulted in reduced pro-coagulant of GPVI-deficient platelets at day 7, while alterations at later time points were only following CRP stimulation (**Supplemental Figure 2B**). In addition to the absence of GPVI on the surface of GPVI-ablated mice (control experiment), we observed elevated GPIbα exposure on the membrane of GPVI-deficient platelets at day 14 and day 28 as compared to WT-PPE controls, following the analysis of GP exposure to the surface of platelets (**Figure 2D–E**). Thereby, vWF binding to platelets (via GPIbα) was only reduced in GPVI-deficient mice after CRP stimulation, under naive conditions as well as at different time points after elastase infusion (**Supplemental Figure 3**). The externalization of integrin β_3_ was reduced after CRP stimulation but not in resting or PAR4 activated GPVI-deficient platelets (**Supplemental Figure 2C**). In contrast to elevated GPIbα exposure at day 14 and day 28 to the membrane of GPVI-deficient platelets, we detected unaltered GP exposure of platelets in naive mice. However, GPVI was absent in GPVI knock-out mice at all time points and under naive conditions, confirming genetic deletion of GPVI in these mice (**Figure 2D, Supplemental Figure 2D**).

**Figure 2.**
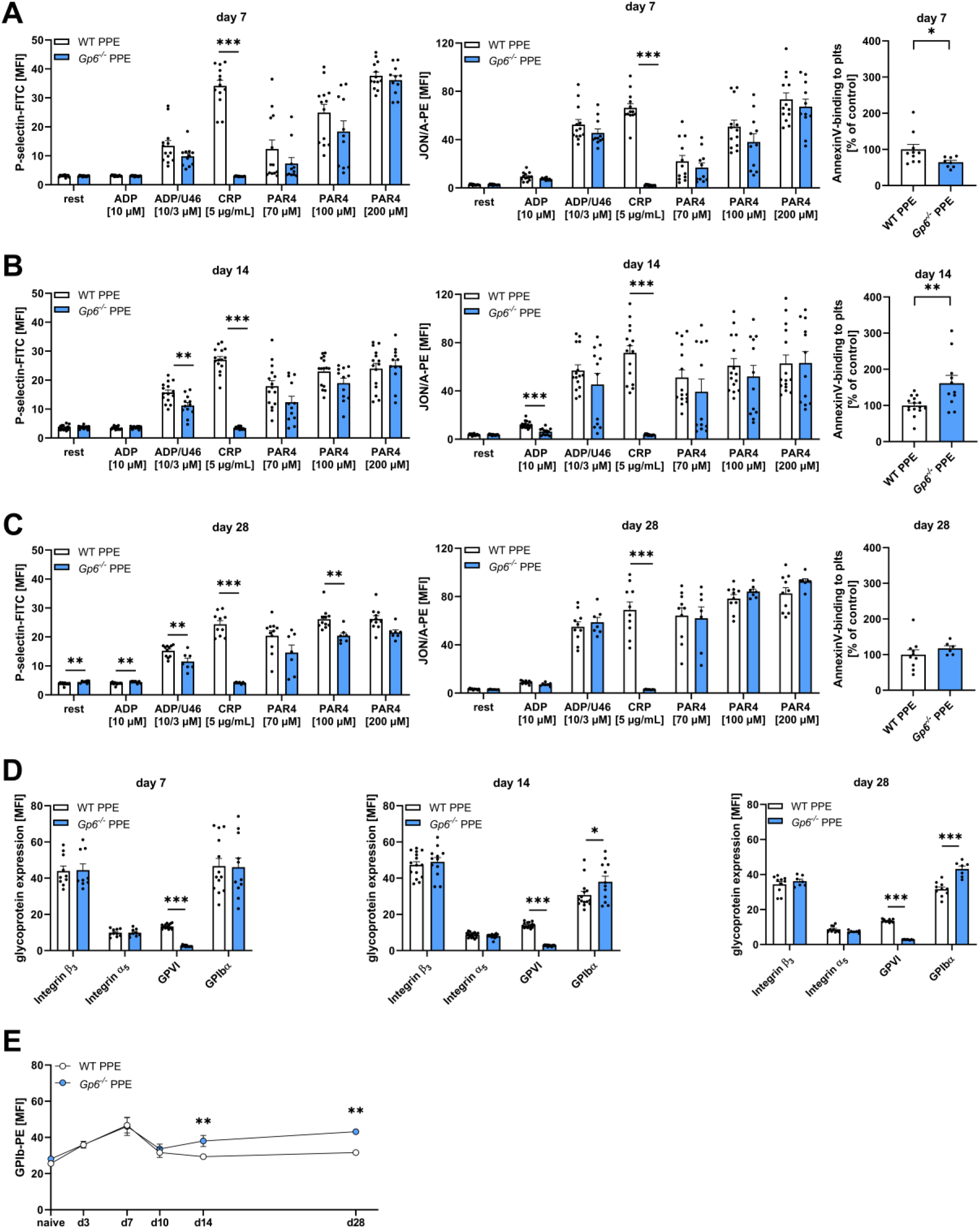
Important role of platelet GPVI in mediating platelet activation and pro-coagulant activity in AAA. (**A–D**) Washed murine whole blood of PPE operated WT and *Gp6^-/-^* mice was analysed via flow cytometry at days 7, 14 and 28 post surgery. (**A–C**) Platelet degranulation (P-selectin-FITC), active integrin α_IIb_β_3_ (JON/A-PE) and PS-exposure of platelets (Annexin V-Cy5) were determined by flow cytometry. Platelets were stimulated with indicated agonists (*Gp6^-/-^* (n=11–15) and WT (n=7–12)). (**D**) Platelet surface exposure of integrin β_3_, integrin α5, GPVI and GPIbα under resting conditions, analysed by flow cytometry (WT (n=7–12) and *Gp6^-/-^* (n=10–15)). (**E**) GPIbα exposure on platelets at day 0 (naive), 3, 7, 10, 14 and 28 after PPE surgery (WT (n=6–12) and *Gp6^-/-^* (n=5–15)). Data are represented as mean values ± SEM. Statistical analysis was performed using (**A–D**) a multiple t-test and (**E**) a two-way ANOVA with a Sidak’s multiple comparisons post-hoc test; *p < 0.05, **p < 0.01, ***p < 0.001. ADP = Adenosine diphosphate; CRP = Collagen-related peptide; MFI, Mean fluorescence intensity; PAR4 = Protease-activated receptor 4 activating peptide; PPE = Porcine pancreatic elastase (infusion); PS = Phosphatidylserine; U46619 (U46) = Thromboxane A_2_ analogue.

### 3.3 Genetic deletion of platelet GPVI impairs inflammation by affecting neutrophil activation and infiltration

To determine the impact of GPVI on platelet-mediated inflammation in experimental AAA, we analysed platelet-leukocyte conjugates in the circulation of PPE mice at different time points. As shown in figure 3A, we detected reduced platelet-leukocyte conjugates at early time point of day 7 (**Figure 3A–B**). In contrast, no differences were observed in the extent of different leukocytes-platelet conjugate formation in naive mice and after 14 days of elastase infusion (**Supplemental Figure 4**). A detailed analysis revealed that reduced conjugates of platelets with CD45 positive cells were preferentially of neutrophil origin based on the reduced number of Ly6G-platelet conjugates in GPVI-deficient mice compared to controls at day 7 (**Figure 3B**) which was not abserved at other time points after elastase infusion or in naive mice (**Supplemental Figure 4C**). The reduced number of neutrophil-platelet conjugates was paralleled by reduced migration of neutrophils and platelets into the aortic wall of GPVI deficient AAA mice (**Figure 3C–D**). In contrast, the migration of macrophages was unaltered (**Figure 3E**). In line with reduced neutrophil presence in the aortic wall, the plasma level of neutrophil elastase as an indicator of neutrophil activation/degranulation was reduced in GPVI-deficient mice (**Figure 3F**). Moreover, the analysis of plasma levels of different pro-inflammatory cytokines at early time point of 7 days post PPE revealed that the acute phase cytokines such as IL1β IL6 and IFN-γ were partly reduced, while anti-inflammatory IL10 levels showed a tendency of increase (**Figure 3G**). This might be due to the fact that thrombo-inflammatory interactions between GPVI activated platelets and leukocytes may foster inflammatory release from the latter; however, genetic ablation of platelet GPVI may not entirely block the release of these mediators.

**Figure 3.**
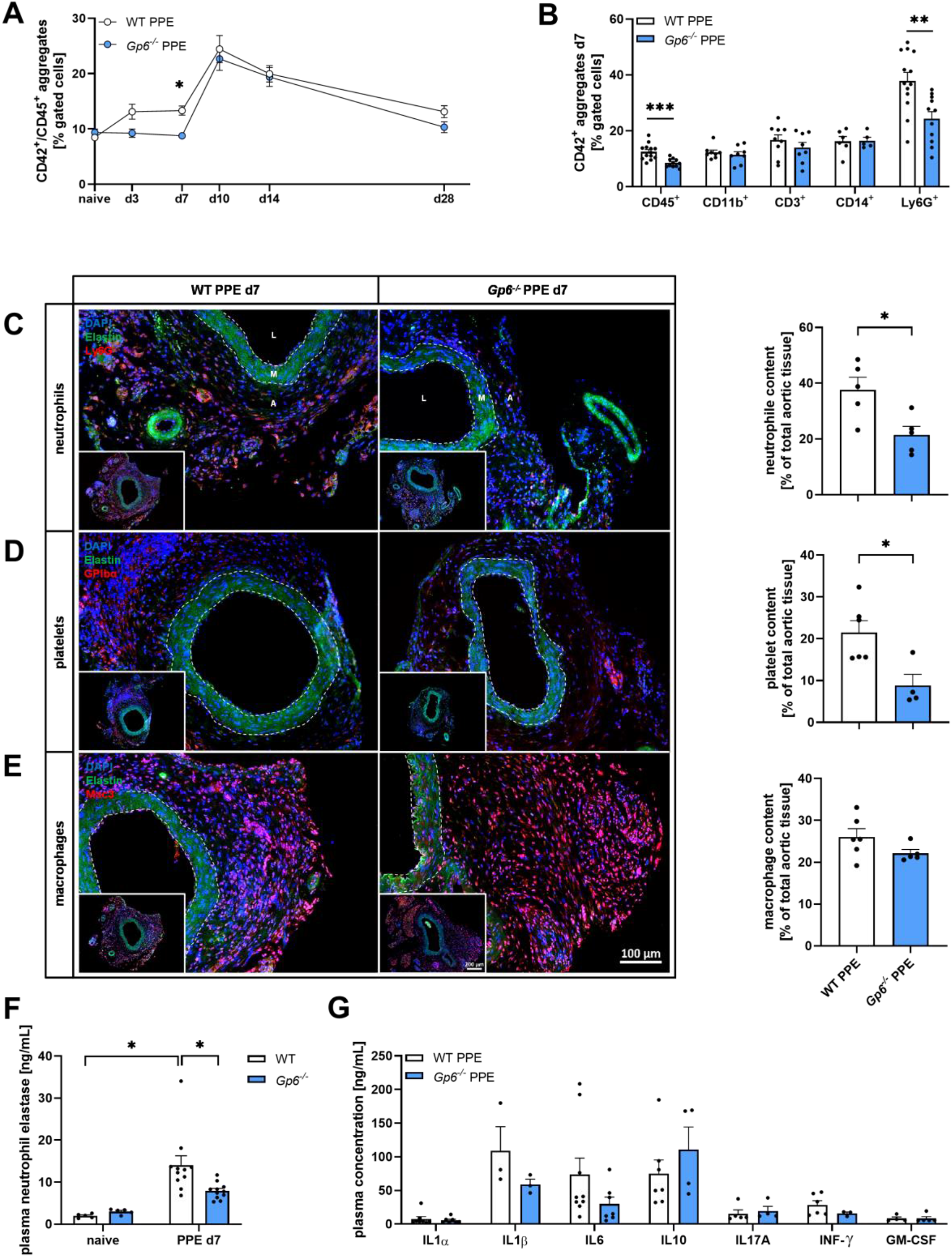
Genetic deletion of platelet GPVI impairs inflammation by affecting neutrophil activation and infiltration. (**A**) Platelet-leukocyte aggregate formation was analysed in whole blood samples of PPE operated WT (n=6–15) and *Gp6^-/-^* (n=6–12) mice at day 0 (naive), 3, 7, 14 and 28 post surgery. Aggregate formation was determined as double positive events for CD42b (platelet maker, GPIbα) and CD45 (leukocyte marker) via flow cytometry. (**B**) Aggregate formation of platelets with different leukocyte subtypes in WT (n=6–14) and *Gp6^-/-^* (n=5–11) PPE mice at day 7 post surgery, analysed via flow cytometry. (**C–E**) Representative immunofluorescence images and quantification of neutrophil, platelet and macrophage infiltration into the aneurysm segment of PPE operated WT (n=5–6) and *Gp6^-/-^* (n=4–5) mice at day 7. Aortic tissue was specifically stained for (**C**) neutrophils (anti-Ly6G/Cy3; red), (**D**) platelets (anti-GPIbα/Cy5; red) or (**E**) macrophages (anti-Mac3/Cy5; red). Elastin autofluorescence (green) is shown (green) and nuclei were stained with DAPI (blue). Scale bar: 100 µm and 200 µm (overview). (**F**) Neutrophil elastase plasma concentration at day 7 post surgery in WT (n=11) and *Gp6^-/-^* (n=11) PPE mice, estimated by ELISA. Naive WT (n=5) and *Gp6^-/-^* (n=5) mice served as controls. (**G**) Plasma concentration of inflammatory cytokines in WT (n=3–8) and *Gp6^-/-^* (n=3–7) PPE mice at day 7 post surgery, determined by multiplex analysis using cytometric bead array. Data are represented as mean values ± SEM. Statistical analysis was performed using (**A**) a two-way ANOVA with a Sidak’s multiple comparisons post-hoc test, (**B** and **G**) a multiple t-test, (**C–E**) an unpaired student’s t-test and (**F**) a two-way ANOVA with a Tukey’s multiple comparisons post-hoc test *p < 0.05, **p < 0.01, ***p < 0.001. A = Adventitia; GM-CSF = Granulocyte-macrophage colony-stimulating factor; IFN = Interferon; IL = Interleukin; L= Lumen; M = Media; OPN = Osteopontin; PPE = Porcine pancreatic elastase (infusion).

### 3.4 Early influence of platelet GPVI in aortic wall remodelling by MMP activation and osteopontin deposition

Next, we investigated whether GPVI affects aortic wall remodelling in AAA. First, we analysed early events such as the presence of matrix metalloproteinases (MMPs) 2 and MMP9 or osteopontin in the plasma of control WT-PPE mice and found increased MMP2 at day 7, high MMP9 at day 14 and increased osteopontin plasma levels at day 7 to day 14 post PPE induction (**Supplemental Figure 5**). Next, we determined MMP and osteopontin plasma levels in GPVI-deficient mice at day 7 after elastase infusion. We detected significantly and strongly reduced plasma levels of MMP2 and osteopontin in the plasma of GPVI-deficient mice while a trend of decrease was observed in MMP9 plasma levels (**Figure 4A–C**). Reduced osteopontin plasma levels were paralleled by reduced osteopontin expression in the aortic wall of GPVI-deficient mice compared to WT-PPE controls (**Figure 4D**). Although, the α-SMA content as marker for the phenotypic switch of fibroblasts to myofibroblasts was unaltered at this early time point, the presence of active caspase-3 positive cells as an indicator of apoptosis was significantly reduced within the aortic tissue of GPVI-deficient mice (**Figure 4E–F**).

**Figure 4.**
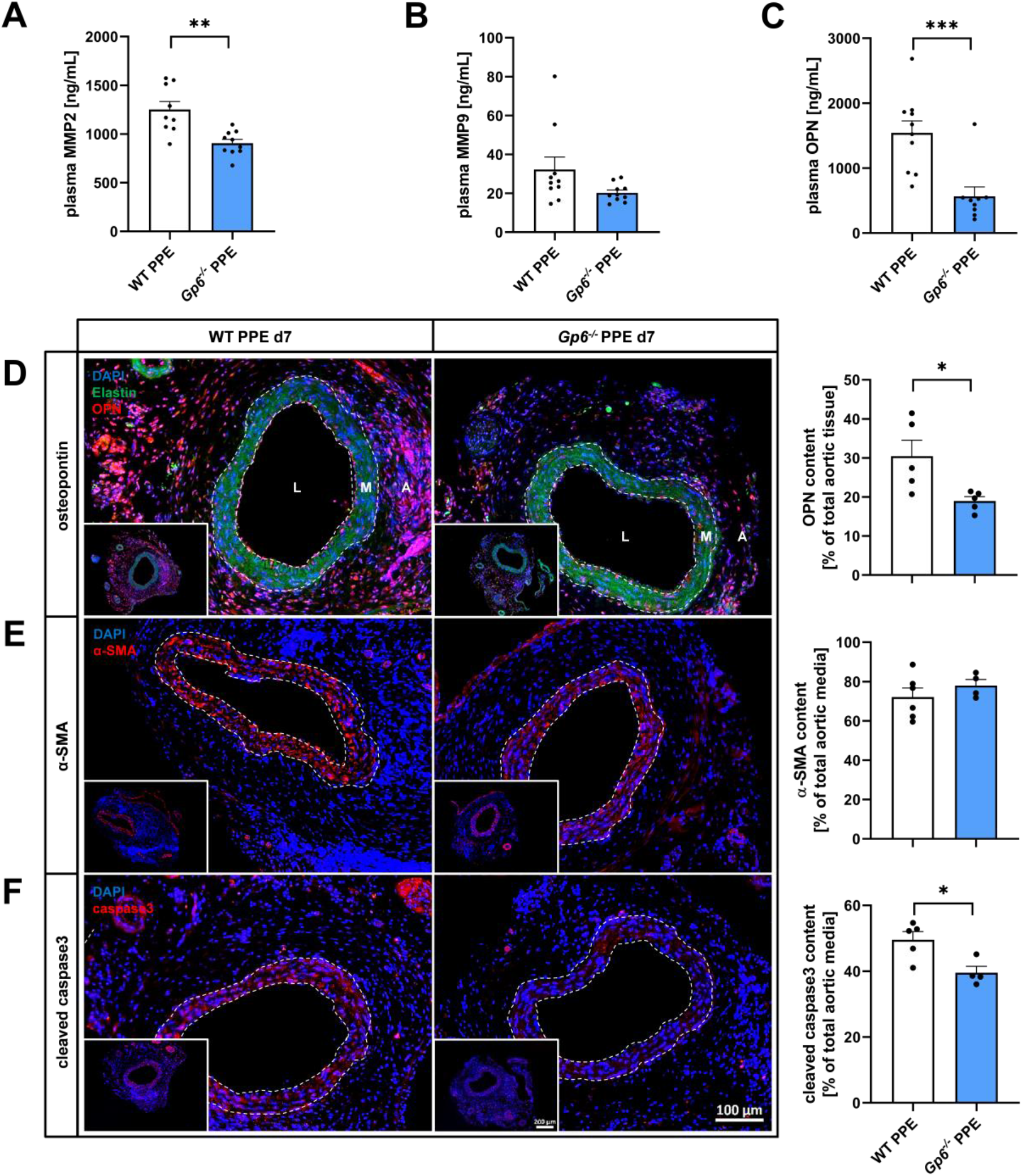
Early interference of GPVI in aortic wall remodelling by MMP activation and osteopontin deposition. (**A–C**) Plasma concentration of (**A**) MMP2, (**B**) MMP9 and (**C**) OPN in PPE operated WT (n=9–11) and *Gp6^-/-^* (n=9–10) mice at day 7 post surgery, determined via ELISA. (**D–F**) Representative immunofluorescence images and quantification of OPN, α-SMA and caspase3 within the aneurysm segment of WT (n=5–6) and *Gp6^-/-^* (n=4–5) PPE mice at day 7. Aortic tissue was specifically stained for (**D**) OPN (anti-OPN/Cy3; red), (**E**) α-SMA (anti-α-SMA/Cy3; red) or (**F**) active caspase3 (anti-cleaved caspase3/Cy3; red). Elastin autofluorescence (green) is shown and nuclei were stained with DAPI (blue). Scale bar: 100 µm and 200 µm (overview). Data are represented as mean values ± SEM. Statistical analysis was performed using (**A–F**) an unpaired student’s t-test; *p < 0.05, **p < 0.01, ***p < 0.001. MMP = Matrix metalloproteinase; OPN = Osteopontin; PPE = Porcine pancreatic elastase (infusion); SMA = Smooth muscle actin.

### 3.5 Modulation of MMP9 release and smooth muscle cell numbers at 14 days after AAA induction

Aortic wall remodelling in AAA was also analysed at day 14 after elastase infusion. At this time point, no differences in MMP2 plasma levels were detected. However, MMP9 and osteopontin plasma levels were significantly reduced in GPVI-deficient mice (**Figure 5A–C**). Reduced osteopontin plasma levels were again paralleled by reduced expression of osteopontin in the aortic wall of GPVI-deficient mice (**Figure 5D**). In addition, we detected significantly upregulated percentage of α-SMA positive cells in the aortic media but comparable caspase-3 positive cells in the aortic wall of GPVI-deficient mice (**Figure 5E–F**).

**Figure 5.**
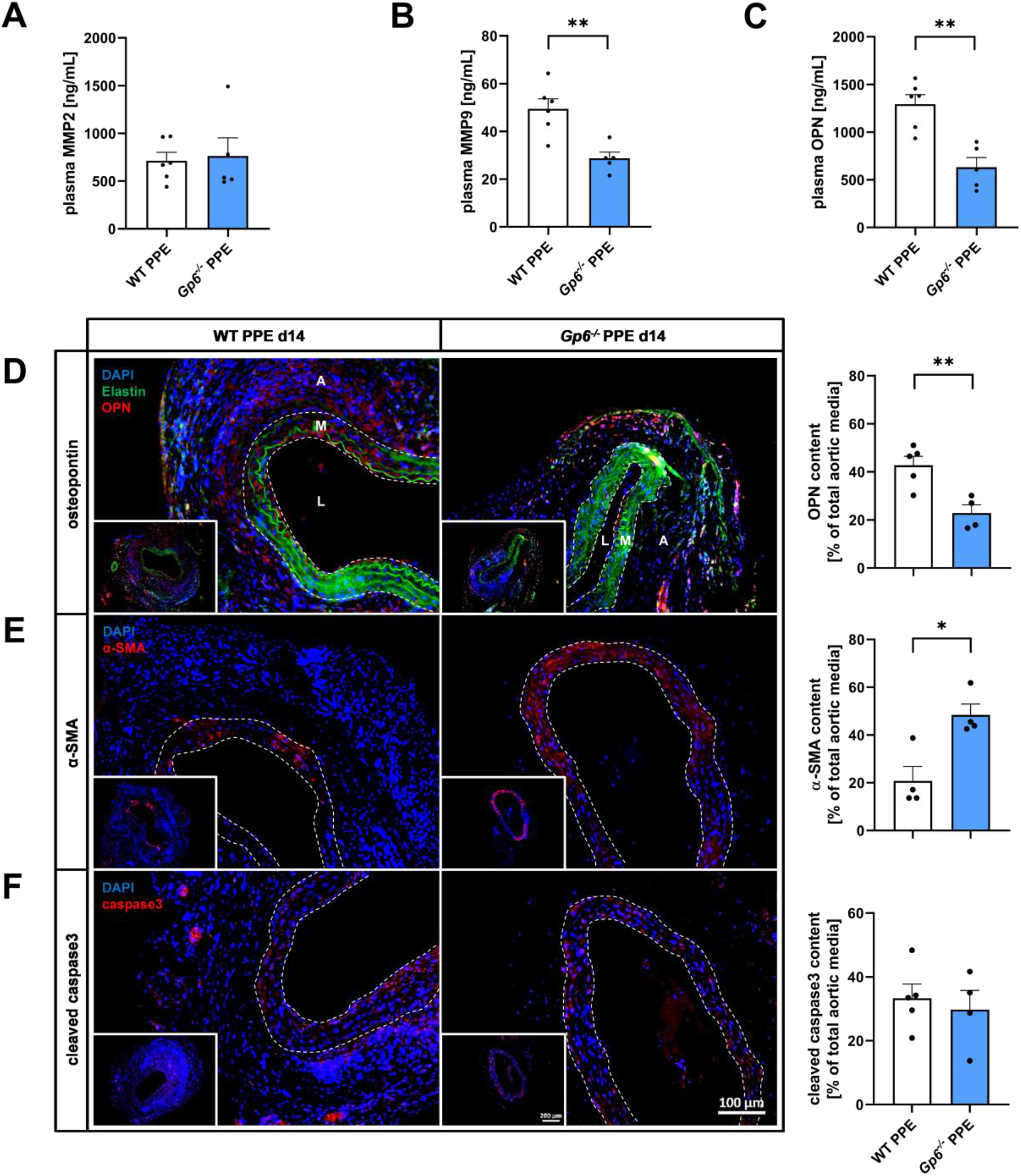
Modulation of MMP9 release and smooth muscle cell numbers at 14 days after AAA induction. (**A–C**) Plasma concentration of (**A**) MMP2, (**B**) MMP9 and (**C**) OPN in PPE operated WT (n=6) and *Gp6^-/-^* (n=5) mice at day 14 post surgery, analysed via ELISA. (**D–F**) Representative immunofluorescence images and quantification of OPN, α-SMA and caspase3 within the aneurysm segment of WT (n=4–5) and *Gp6^-/-^* (n=4) PPE mice at day 14. Aortic tissue was specifically stained for (**D**) OPN (anti-OPN/Cy3; red), (**E**) α-SMA (anti-α-SMA/Cy3; red) or (**F**) active caspase3 (anti-cleaved caspase3/Cy3; red). Elastin autofluorescence (green) is shown and nuclei were stained with DAPI (blue). Scale bar: 100 µm and 200 µm (overview). Data are represented as mean values ± SEM. Statistical analysis was performed using (**A–F**) an unpaired student’s t-test; *p < 0.05, **p < 0.01, ***p < 0.001. MMP = Matrix metalloproteinase; OPN = Osteopontin; PPE = Porcine pancreatic elastase (infusion); SMA = Smooth muscle actin.

### 3.6 Platelet GPVI significantly contributes to aortic wall remodelling in AAA

Differences in MMP plasma levels, osteopontin content and α-SMA positive cells in GPVI-deficient PPE mice resulted in improved aortic wall remodelling 28 days after elastase infusion as shown by increased intima/media thickness, reduced elastin fragmentation and elevated elastin content (**Figure 6A–C**). Improved aortic wall remodelling in GPVI-deficient mice was reflected by a strong correlation (determined by the Spearman’s correlation coefficient) of intima/media thickness, as well as elastin fragmentation to aneurysm diameter progression at day 28 post elastase infusion (**Figure 6D–E**). In contrast to the elastin content, we did not observe major changes in the average collagen content in GPVI-deficient PPE mice compared to WT-PPE controls (**Figure 6F**). However, a detailed analysis of the collagen of the aortic wall revealed an altered collagen composition based on a significantly elevated tight collagen and reduced fine collagen in mice lacking GPVI (**Figure 6G**). In control experiments, we provided evidence that no differences were observed in aortic intima/media thickness, elastin fragmentation and collagen composition in naive mice from either groups (**Supplemental figure 6**).

**Figure 6.**
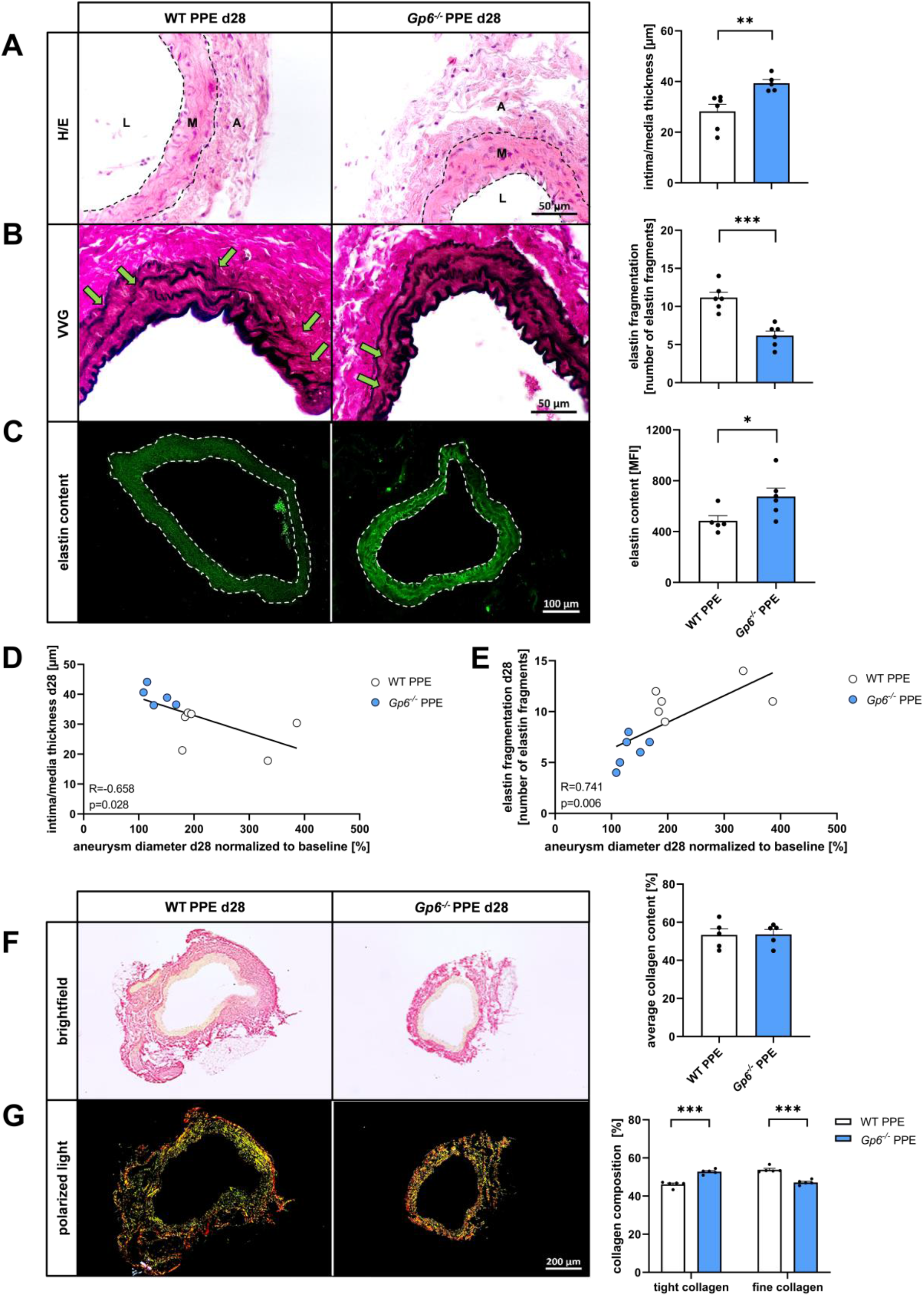
Platelet GPVI significantly contributes to aortic wall remodelling in AAA. (**A– C**) Representative images and quantification of aortic tissue from PPE operated WT and *Gp6^-/-^* mice at day 28 post surgery. (**A**) The intima/media thickness was analysed using H/E staining of WT (n=6) and *Gp6^-/-^* (n=5) PPE mice. Scale bar: 50 µm. (**B**) Quantification of elastin fragmentation within the aortic media of WT (n=6) and *Gp6^-/-^* (n=6) PPE mice via VVG staining. Scale bar: 50 µm. (**C**) Elastin content of WT (n=4) and *Gp6^-/-^* (n=6) PPE mice by analyzing the elastin autofluorescence within the aortic media. Scale bar: 100 µm. (**D** and **E**) Spearman’s correlation between (**D**) intima/media thickness, respectively (**E**) elastin fragmentation and the aneurysm diameter at day 28 post PPE. (**F** and **G**) Collagen content and composition in aortic tissue of PPE operated WT (n=5) and *Gp6^-/-^*(n=5) at day 28. (**F**) Representative brightfield images and quantification of the average collagen content using Picrosirius red staining. (**G**) Representative images and quantification of the aortic collagen composition using polarized light. Aortic tissue was stained using Picrosirius red. Scale bar: 200 µm. Data are represented as mean values ± SEM. Statistical analysis was performed using (**A–C** and **F**) an unpaired student’s t-test and (**G**) a two-way ANOVA with a Sidak’s multiple comparisons post-hoc test; *p < 0.05, **p < 0.01, ***p < 0.001. A = Adventitia; H/E = Hematoxylin-eosin; L = Lumen; M = Media; MFI = Mean fluorescence intensity; PPE = Porcine pancreatic elastase (infusion); VVG = Verhoeff van Gieson.

### 3.7 Elevated plasma levels of soluble GPVI and enhanced deposition of the GPVI ligand fibrin in the intraluminal thrombus and in the plasma of AAA patients

To study the relevance of platelet GPVI in AAA pathology among patients, we first determined levels of soluble (s) GPVI in the plasma and surface expression of GPVI at the platelet surface of AAA patients. We detected increased sGPVI plasma levels while the abundance of GPVI at the platelet surface was not significantly different compared to healthy controls (**Figure 7A and B**). Moreover, the platelet content within the ILT of AAA patients was reduced compared to arterial thrombi of different origin, isolated from patients that underwent thrombectomy (**Figure 7D**). In contrast, the content of fibrin, an identified GPVI ligand, important for GPVI mediated thrombus stabilization (47), was highly elevated in the ILT of AAA patients compared to arterial thrombi from thrombectomy patients (**Figure 7E**). Increased fibrin deposition in the ILT was paralleled by elevated plasma levels of fibrin in AAA patients compared to controls (**Figure 7C**). Furthermore, thrombus formation using peripheral blood samples in the *ex vivo* flow chamber system revealed no differences in thrombus formation or volume on collagen under dynamic arterial flow conditions when we compared blood from controls and AAA patients (**Figure 7F**). This might be due to the existing localised thrombo-inflammatory environment in the ILT contributing to the pathological progression of AAA. Clinical parameter of the analysed AAA and thrombactomy patient cohort are summarized in Supplemental Table 1.

**Figure 7.**
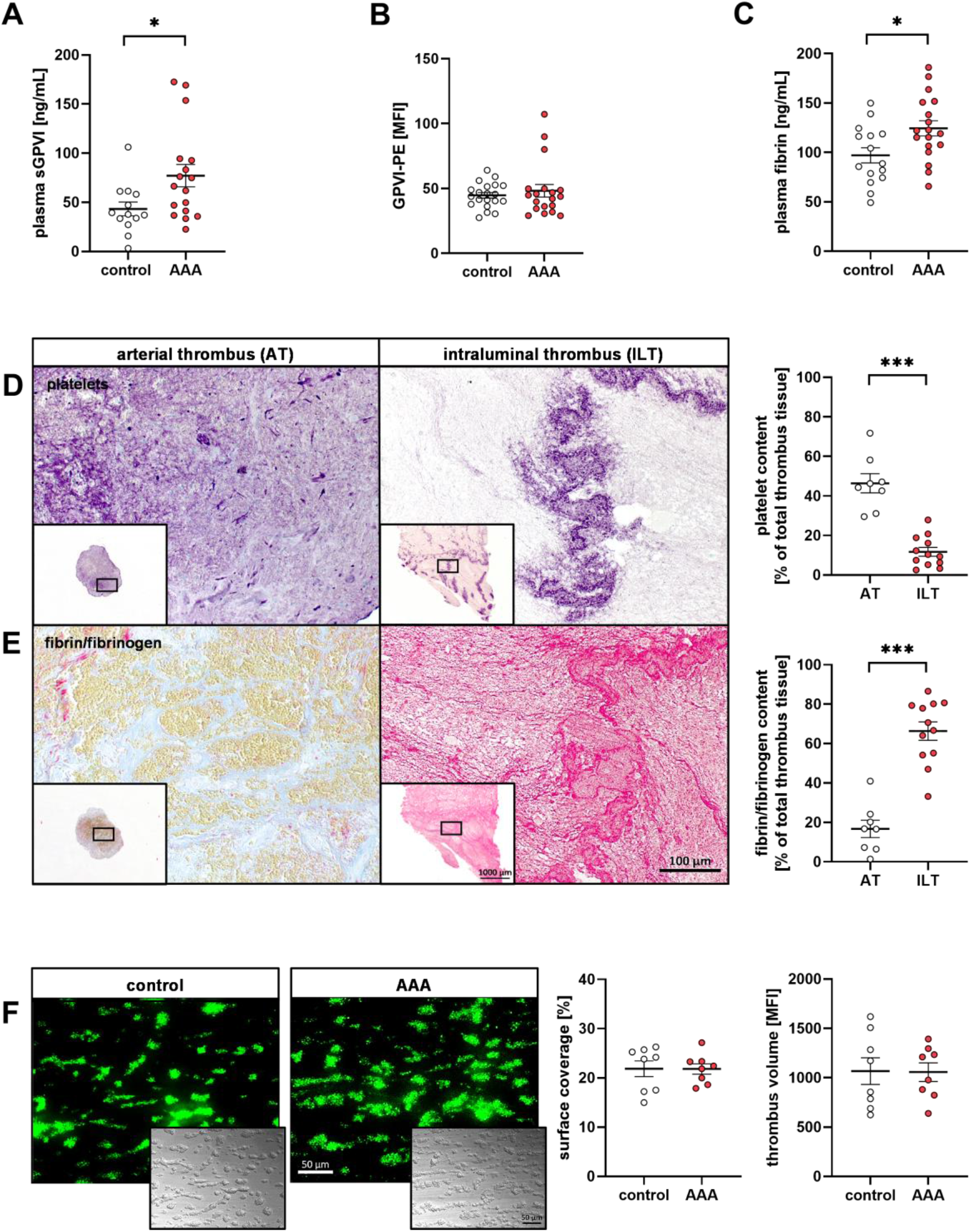
Elevated plasma levels of soluble GPVI and enhanced deposition of the GPVI ligand fibrin in the intraluminal thrombus and in the plasma of AAA patients. (**A**) Plasma concentration of sGPVI in samples of AAA (n=18), analysed via ELISA. Samples of healthy volunteers served as controls (n=13). (**B**) Platelet surface GPVI exposure of AAA (n=19) patients, determined in whole blood samples by flow cytometry. Healthy volunteers served as controls (n=20). (**C**) Fibrin plasma concentration in samples of AAA (n=18), analysed via ELISA. Samples of healthy volunteers served as controls (n=15). (**D** and **E**) Representative images and quantification of intraluminal thrombus (ILT) tissue samples from AAA patients (n=12). Arterial thrombus (AT) tissue from thrombectomy patients (n=8) served as control. (**D**) Thrombus sections were immunohistochemically stained specifically for platelets (anti-GPIbα). (**E**) Histological MSB staining of thrombus sections to visualize fibrin depositions. Scale bar: 100 µm and 1000 µm (overview). **(F)** Representative fluorescence images and quantification of *ex vivo* flow chamber experiments. Whole blood samples of AAA (n=8) patients was perfused over a collagen matrix (200 µg/mL) under an arterial shear rate of 1,700 s^-1^. Whole blood samples of healthy volunteers served as controls (n=8). Scale bar: 50 µm. Data are represented as mean values ± SEM. Statistical analysis was performed using (**A–F**) an unpaired student’s t-test; *p < 0.05, ***p < 0.001. AAA = Abdominal aortic aneurysm; AT = arterial thrombus; ILT = Intraluminal thrombus; MSB = Martius scarlet blue; sGPVI = soluble GPVI.

## 4. Discussion

In this study, we found that genetic deletion of the major platelet collagen receptor GPVI protects mice against aortic diameter expansion in experimental AAA. These protective effects are attributed to reduced inflammation due to decreased platelet-neutrophil aggregate formation, neutrophil activation and infiltration into the aortic wall. In addition, GPVI promotes remodelling of the aortic wall as reflected by reduced MMP2/9 and OPN plasma levels but elevated content of α-SMA in the aortic tissue accompanied by reduced cell apoptosis. Consequently, enhanced intima/media thickness and elevated elastin content were observed in GPVI-deficient PPE mice, leading to significantly reduced aortic diameter expansion and reduced aneurysm incidence. Importantly, AAA patients showed elevated plasma levels of sGPVI and fibrin, accompanied by massive fibrin accumulation within the ILT, suggesting that platelet GPVI might be a crucial mediator in thrombo-inflammatory processes. Moreover, sGPVI could serve as a potential biomarker for AAA development as in other cardiovascular atherothrombotic diseases. In addition, binding of platelet GPVI to fibrin might stabilize ILT formation and progression in the aorta of AAA patients

AAA is an atherosclerotic-related disease with multifactorial causes. Characteristic hallmarks are chronic inflammation and degradation of the ECM within the abdominal aortic wall, often accompanied by the formation of an ILT in AAA patients. Abnormal platelet activation has been described for many atherosclerotic, cardiovascular diseases (CVDs) leading to an increased risk for thromboembolic events (29). In AAA, elevated platelet activation has been described as a major pathological feature in experimental mice (PPE mouse model) and in AAA patients. Specifically, platelet hyperresponsiveness/hyperreactivity in both mice and patients with AAA have been described whereby platelet GPVI-collagen/fibrinogen/fibrin interactions may trigger AAA initiation and progression (19). In addtion, another study provided evidence for enhanced activation after stimulation of platelets with thrombin and thromboxane (16). In the present study, we detected reduced platelet activation and pro-coagulant activity in response to PAR4 peptide (Annexin-V binding, P-selectin exposure) that might be responsible –at least in part– for reduced aortic diameter expansion in GPVI-deficient mice. Within the whole the 28-day observation period, reduced pro-coagulant activity of GPVI-deficient platelets was detected in response to GPVI stimulation but interestingly also in resting and PAR4 stimulated platelets at early time points (day 7). It has already been shown that GPVI activation is able to directly support thombin-induced platelet activation. Interestingly, this process is independent of Syk and Src kinases (30). In a more indirect way, GPVI is able to enhance thrombin mediated platelet activation by triggering PS exposure to provide a pro-coagulant surface at the platelet membrane leading to thombin generation (31, 32).

Reduced pro-coagulant activity of GPVI-deficient platelets might result from decreased platelet activation. Elevated thrombin plasma levels in AAA patients might occur as a consequence of the enhanced pro-coagulant activity of platelets which is a characteristic feature of AAA (33, 34). Thus, we believe that –together with reduced platelet activation- and procoagulant activity of GPVI-deficient platelets might account for reduced aortic diameter expansion in these mice. Furthermore, our previously mentioned study provided strong evidence for platelets to trigger osteopontin expression and release from macrophages via GPVI, indicating that osteopontin not only plays a role in inflammatory and remodelling processes in AAA but might modulate recruitment of platelets to the aortic wall and the ILT. Thus, it is plausible that the reduced osteopontin protein levels in the plasma and the aortic wall of GPVI-deficient mice resulted in reduced accumulation/infiltration of platelets into the aortic wall at day 7 after PPE induction. (19).

To date, only a small number of studies that mainly utilized the Ang-II infusion model of experimental AAA have analysed the role of platelets. Blocking of the platelet ADP receptor P2Y12 by clopidogrel was shown to prevent AAA formation by protecting the elastic lamina of the mouse aorta and by reduced inflammation and MMP2 expression (35). Recently, we have shown that platelet depletion by antibody treatment protects mice from aortic diameter expansion (19), suggesting a dominant role of platelets in AngII- and elastase-induced AAA formation in mice. Other investigators have explored the role of platelets in the (e)PPE mouse model. Here, the authors detected elevated platelet activation in mice and patients (16) with a role of P-selectin because P-selectin knock-out mice showed attenuated AAA (14).

Platelet GPVI has been previously shown to affect inflammatory response and vascular integrity. Boilard and colleagues identified GPVI to trigger microparticle production, amplifying inflammation in arthritis (36). Moreover, GPVI has been shown to contribute to local host defence during pneumonia driven sepsis by triggering leukocyte activation (37). During cutaneous inflammation, GPVI promotes the inflammatory response of macrophages (38). Importantly, platelet recruitment to the site of inflammation is mediated – at least in part – by GPVI as shown in a model of glomerulonephritis (39).

Beside the impact of platelet GPVI in inflammation, an important role of this collagen receptor has been shown in different cerebro- and cardiovascular diseases both in patients and in experimental rodent models (24, 40–42). Furthermore, GPVI is not only involved in different cellular processes in cerebro- and cardiovascular diseases but also represents a useful biomarker for the early detection of these diseases and has great potential for strategizing an efficacious anti-thrombotic therapy (40). To date, the translation of experimental findings into therapeutic implications for AAA has benn challenging. Several trials for different drugs and therapeutic strategies have failed to proof a clinical benefit (43–45). There were also different trials that analysed the benefit of anti-platelet therapy (11, 45–47). However, it remains an opon question, which AAA patients could benefit most from anti-platelet therapy. Interestingly, major differences in the efficacy of anti-platelet therapy have been linked to aneurysm size because patients with small-size aneurysms often did not benefit from anti-platelet therapy, while beneficial effects have been demonstrated for patients with large aneurysms (11, 45, 46). Therapeutic benefits of anti-platelet drugs in enlarged AAAs might be due to the fact that these patients develop an ILT that is only rarely present in patients with small AAA (11). Anti-platelet therapy targeting GPVI might be a noval option in the therapy of AAA, since different experimental studies and clinical trials showed beneficial effects by blocking GPVI in acute myocardial infarction or stroke in mice or by the treatment of stroke patients with the GPVI Fab ACT017 (Glenzocimab) (42, 48, 49). The idea of GPVI to represent a new therapeutic target in AAA is emphasized by the results from AAA patients (**Figure 7**) demonstrating elevated levels of sGPVI and fibrin in the plasma as well as increased fibrin deposition in the ILT of these patients. Fibrin is a GPVI ligand that is able to stabilize thrombus formation under dynamic conditions (50). Furthermore, binding of GPVI to fibrin has been shown to promote procoagulant activity and impacts clot structure suggesting a substantial role of the GPVI-fibrin axis in AAA by increasing thrombin generation and stabilizing the ILT (51, 52).

In conclusion, our current data supports the potential role of GPVI as new anti-thrombotic therapy in AAA. However, interfering with the GPVI-fibrin axis might lead to destabilization of the ILT thus its role in ILT formation and clot structure has to be carefully evaluated.

## Supporting information

Feige et al_Suppl_2023_final

## Acknowledgments

We thank Martina Spelleken and Annika Zimmermann for excellent technical assistance. PPE surgeries were conducted by Julia Odendahl (Department of Cardiology, Pulmonology and Vascular Medicine, University Hospital Duesseldorf; S1-Project TRR259).

## Author contributions

ME designed the study. TF, AB, KJK, JM, MC, JO, EK and IK performed experiments. TF, KJK, MC, and ME analysed and interpreted data. TF and ME wrote the manuscript with all authors providing feedback.

## Data Availability

The original contributions presented in the study are included in the article, further inquiries can be directed to the corresponding author.

