## Supplementary material for "Collagen receptor GPVI-mediated platelet activation and pro-coagulant activity aggravates inflammation and aortic wall remodelling in abdominal aortic aneurysm": Feige et al_Suppl_2023_final

* contributed equally

§ corresponding author

### Supplemental figures

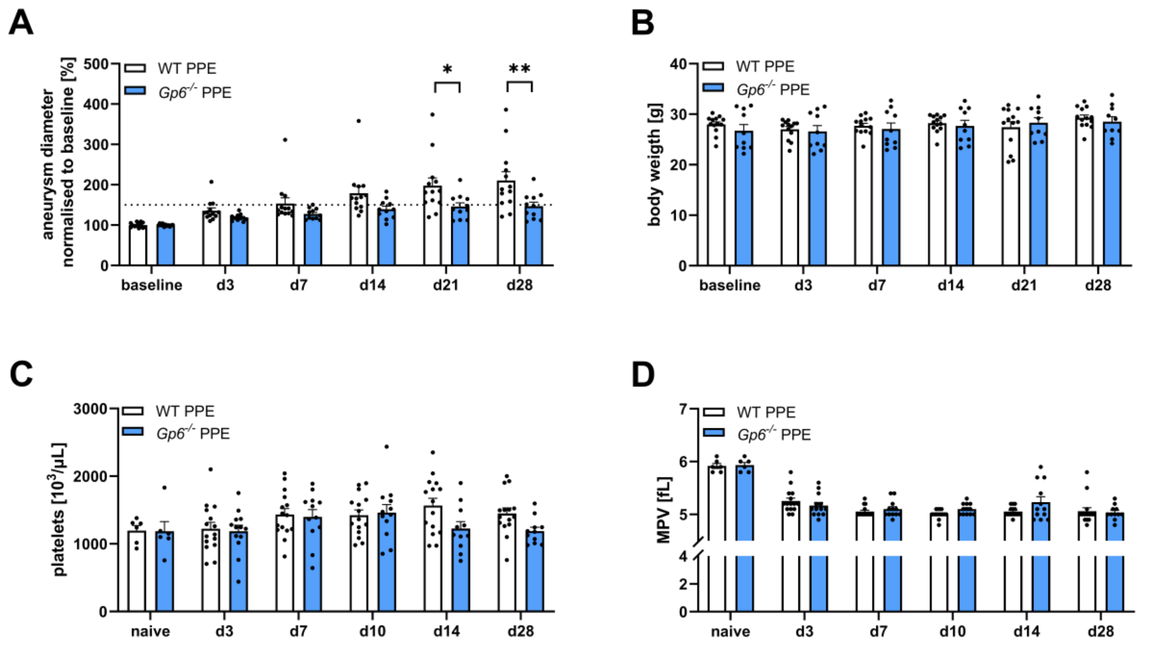

**Figure S1. Aortic diameter, body weight and platelet count of GPVI-deficient mice in PPE-induced AAA progression.** (**A**) Aortic diameter progression of WT (n=13) and *Gp6*$^{-/-}$ (n=11) PPE mice over a time period of 28 days, analysed by ultrasound measurements. Data were normalised to baseline. (**B**) Body weight of PPE operated WT (n=13) and *Gp6*$^{-/-}$ (n=11) mice prior (baseline) and at day 3, 7, 14, 21, 28 post surgery. (**C** and **D**) Platelet count and MPV of naive (WT (n=6) and *Gp6*$^{-/-}$ (n=6)) and PPE operated (WT (n=13) and *Gp6*$^{-/-}$ (n=11) mice at day 7, 14 and 28 post surgery. (Data are represented as mean values ± SEM. Statistical analysis was performed using (**A–D**) a two-way ANOVA with a Sidak's multiple comparisons post-hoc test and (**C** and **D**) an unpaired student's t-test; *p < 0.05, **p < 0.01. AAA = Abdominal aortic aneurysm; MPV = Mean platelet volume.

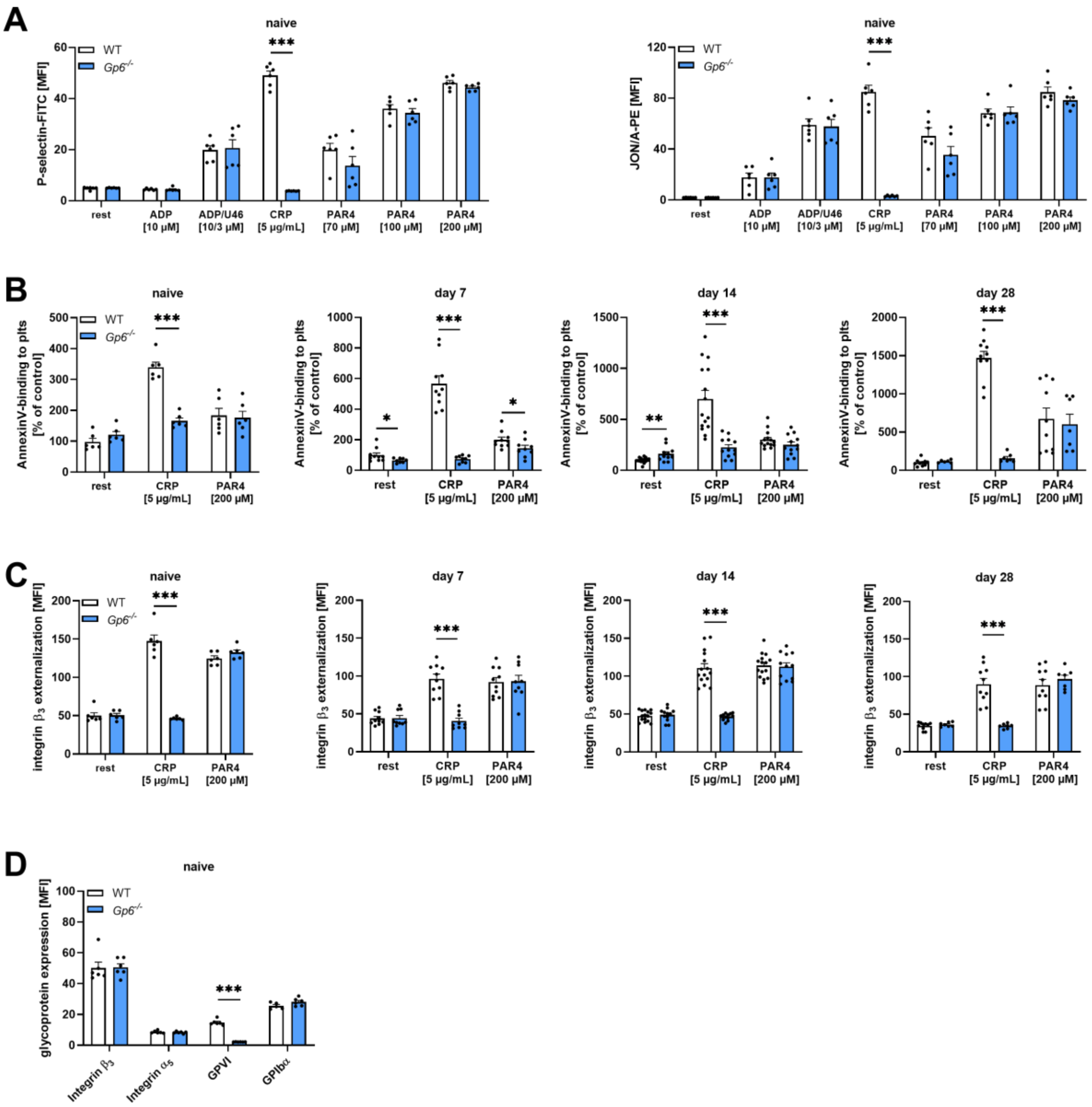


**Figure S2. Integrin β$_3$ externalization and PS exposure of GPVI-deficient mice in PPE-induced AAA progression.** (**A–D**) Washed murine whole blood of naive and PPE operated WT and *Gp6*$^{-/-}$ mice was analysed by flow cytometry. (**A**) Platelet degranulation (P-selectin-FITC) and active integrin α$_{IIb}$β$_3$ (JONA/A-PE) of naive WT (n=6) and *Gp6*$^{-/-}$ (n=6) mice were determined by flow cytometry. Platelets were stimulated with indicated agonists. (**B**) PS-exposure (Annexin V-Cy5) of platelets from naive (WT (n=6) and *Gp6*$^{-/-}$ (n=6) and PPE operated (WT (n=8–14) and *Gp6*$^{-/-}$ (n=9–12)) mice at day 7, 14 and 28 post surgery. Platelets were stimulated with indicated agonists. (**C**) Integrin β$_3$ externalization in naive WT (n=6) and *Gp6*$^{-/-}$ (n=6) and PPE operated (WT (n=10–15) and *Gp6*$^{-/-}$ (n=7–12)) mice at day 7, 14 and 28 post surgery. Platelets were stimulated with indicated agonists. (**D**) Platelet surface expression of integrin β$_3$, integrin α$_5$, GPVI and GPIbα under resting conditions in naive WT (n=6) and *Gp6*$^{-/-}$ (n=6) mice. Data are represented as mean values ± SEM. Statistical analysis was performed using (**A–D**) a multiple t-test; ***p < 0.001. ADP = Adenosine diphosphate; CRP =

Collagen-related peptide; MFI = Mean fluorescence intensity; PAR4 = Protease-activated receptor 4 activating peptide; PS = Phosphatidylserine; PPE = Porcine pancreatic elastase (infusion); U46619 (U46) = Thromboxane $A_2$ analogue.

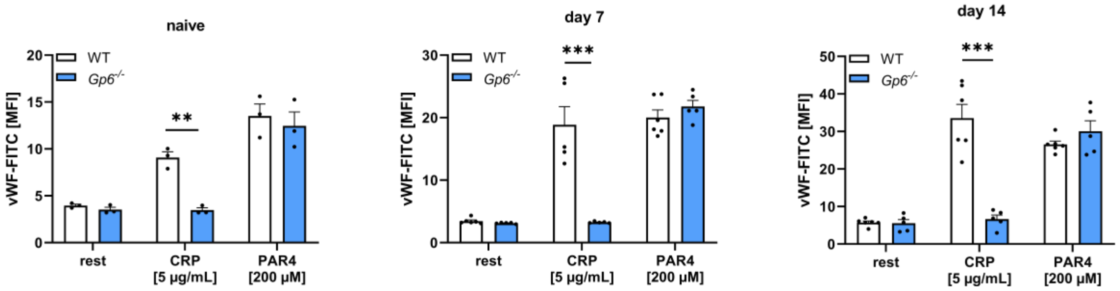


**Figure S3. vWF deposition on the platelet surface of GPVI deficient mice during PPE-induced AAA development.** Washed murine whole blood of naive (unoperated) and PPE operated WT and *Gp6*$^{-/-}$ mice were analysed by flow cytometry. vWF-binding to platelets from naive (WT (n=3) and *Gp6*$^{-/-}$ (n=3)) and PPE operated (WT (n=5-6) and *Gp6*$^{-/-}$ (n=5)) mice at day 7 and 14 post surgery. Platelets were stimulated with indicated agonists. Data are represented as mean values ± SEM. Statistical analysis was performed using a multiple t-test; ***p < 0.001. ADP = Adenosine diphosphate; CRP = Collagen-related peptide; MFI = Mean fluorescence intensity; PAR4 = Protease-activated receptor 4 activating peptide; PPE = Porcine pancreatic elastase (infusion); U46619 (U46) = Thromboxane $A_2$ analogue; vWF = van Willeband factor.

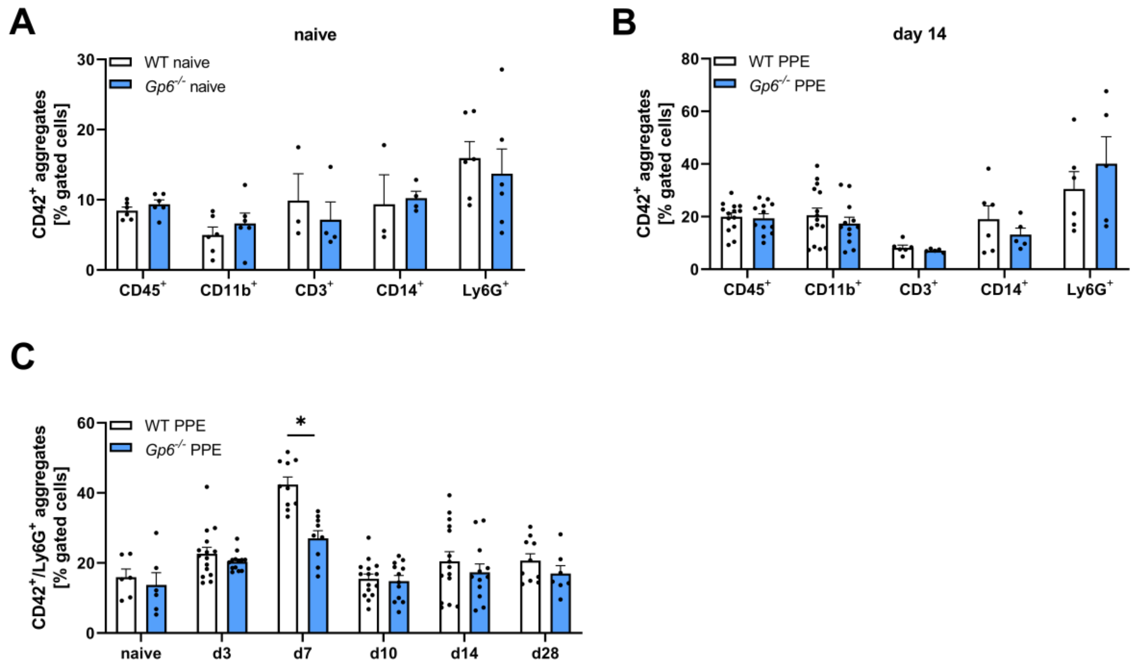


**Figure S4. Platelet-leukocyte aggregates in naive and PPE operated GPVI-deficient mice.** (**A** and **B**) Aggregate formation of platelets with different leukocyte subtypes in (**A**) naive (WT (n=3–6) and *Gp6*$^{-/-}$ (n=3–6)) and (**B**) PPE operated (WT (n=6–14) and *Gp6*$^{-/-}$ (n=5–12)) mice at day 14 post surgery, analysed by flow cytometry. (**C**) Platelet-neutrophil aggregate formation was analysed in PPE operated WT (n=6–15) and *Gp6*$^{-/-}$ (n=6–12) mice at day 0 (naive) 3, 7, 14, 28 post surgery. Aggregate formation was determined as double positive events for CD42b (platelet maker, GPIbα) and CD45 (WBC marker) by flow cytometry. Data are represented as mean values ± SEM. Statistical analysis was performed using (**A** and **B**) a multiple t-test and (**C**) a two-way ANOVA with a Sidak's multiple comparisons post-hoc test *p < 0.05. PPE = Porcine pancreatic elastase (infusion).

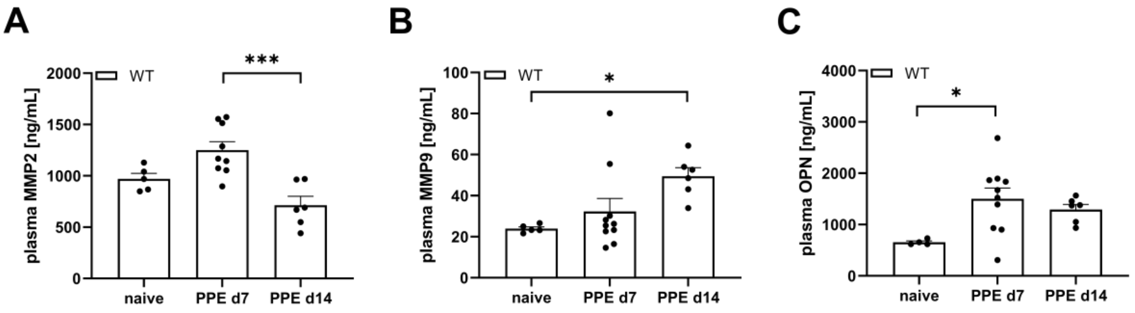


**Figure S5. MMP and OPN plasma level during experimental AAA progression.** (**A–C**) Plasma concentration of (**A**) MMP2, (**B**) MMP9 and (**C**) OPN of PPE-operated WT (n=6–10) mice at day 7 and 14 post surgery, analysed by ELISA. Naive WT (n=4–5) mice served as control. Data are represented as mean values ± SEM. Statistical analysis was performed using (**A–C**) a one-way ANOVA with a Tukey's multiple comparisons post-hoc test; *p < 0.05, ***p < 0.001. MMP = Matrix metalloproteinase; OPN = Osteopontin; PPE = Porcine pancreatic elastase (infusion).

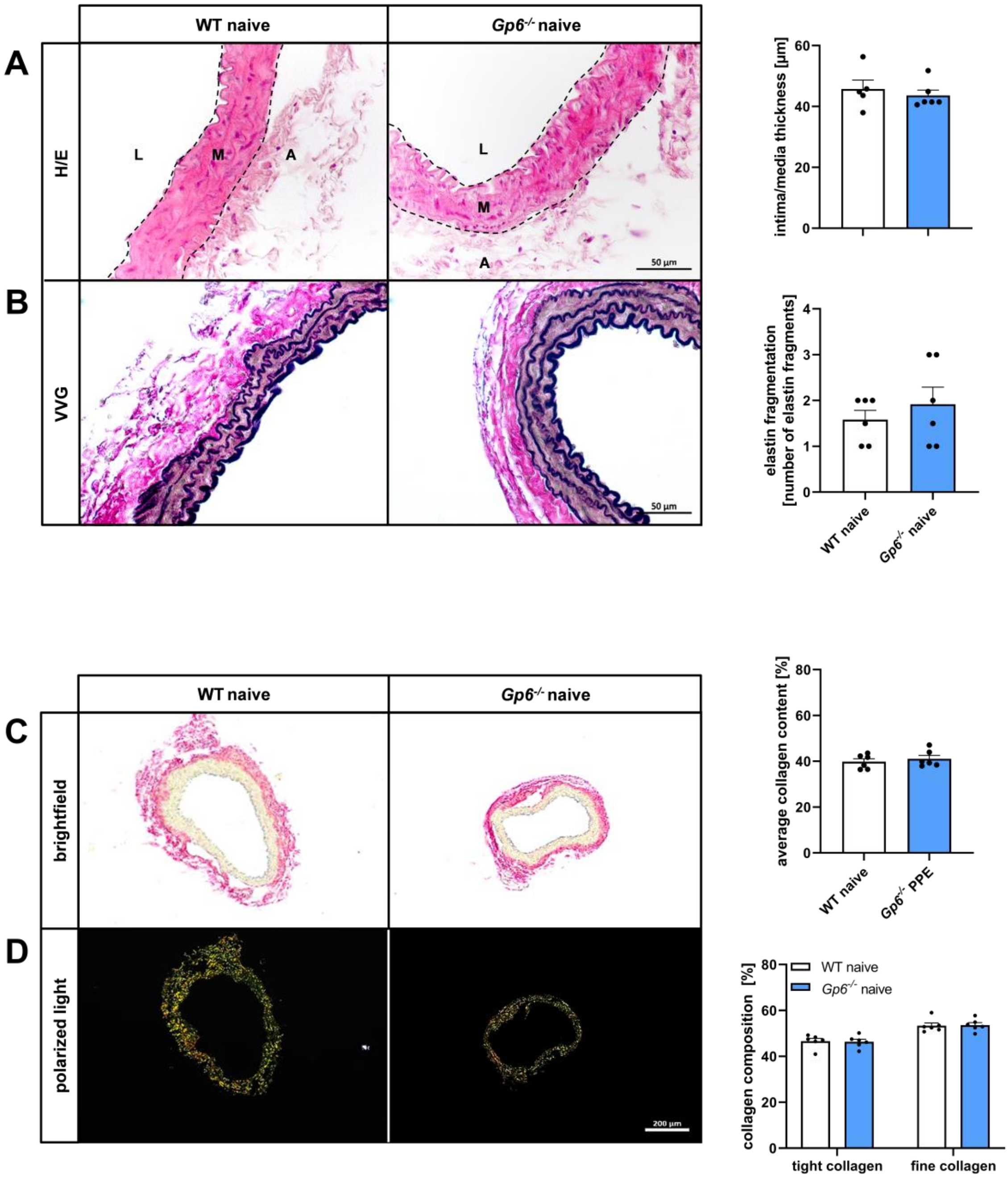


**Figure S6. Aortic intima/media thickness, elastin fragmentation and collagen composition in naive GPVI-deficient mice.** (**A–D**) Representative images and quantification of aortic tissue from naive (unoperated) WT and *Gp6*$^{-/-}$ mice. (**A**) Intima/media thickness was analysed by using H/E staining of naive WT (n=5) and *Gp6*$^{-/-}$ (n=6) mice. Scale bar: 50 µm. (**B**) Quantification of elastin fragmentation within the aortic media of naive WT (n=6) and *Gp6*$^{-/-}$ (n=6) mice by VVG staining. Scale bar: 50 µm. (C) Representative brightfield images and quantification of the average collagen content using Picrosirius red staining. (**G**) Representative images and quantification of the aortic collagen composition using polarized light. Aortic tissue was stained using Picrosirius red. Scale bar: 200 µm. Data are represented as mean values ± SEM. Statistical analysis was performed using (**A–D**) an unpaired student's t-test. A = Adventitia; H/E = Hematoxylin-eosin; L = Lumen; M = Media; PPE = Porcine pancreatic elastase (infusion); VVG = Verhoeff van Gieson.

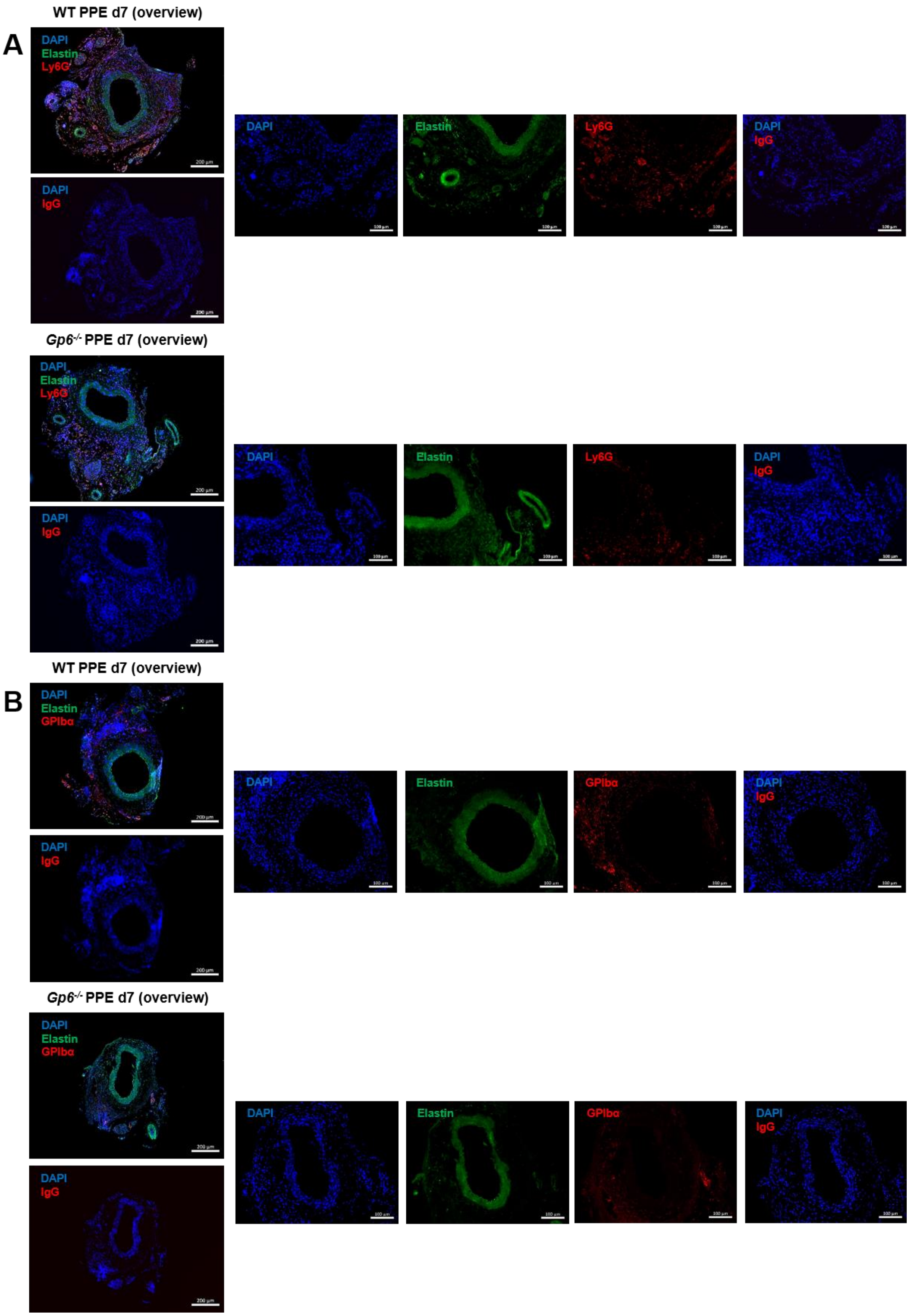
WT PPE d7 (overview)
A
DAPI
Elastin
Ly6G
DAPI
IgG
DAPI
Elastin
Ly6G
DAPI
IgG
Gp6-/- PPE d7 (overview)
DAPI
Elastin
Ly6G
DAPI
IgG
DAPI
Elastin
Ly6G
DAPI
IgG
WT PPE d7 (overview)
B
DAPI
Elastin
GPIbα
DAPI
IgG
DAPI
Elastin
GPIbα
DAPI
IgG
Gp6-/- PPE d7 (overview)
DAPI
Elastin
GPIbα
DAPI
IgG
DAPI
Elastin
GPIbα
DAPI
IgG

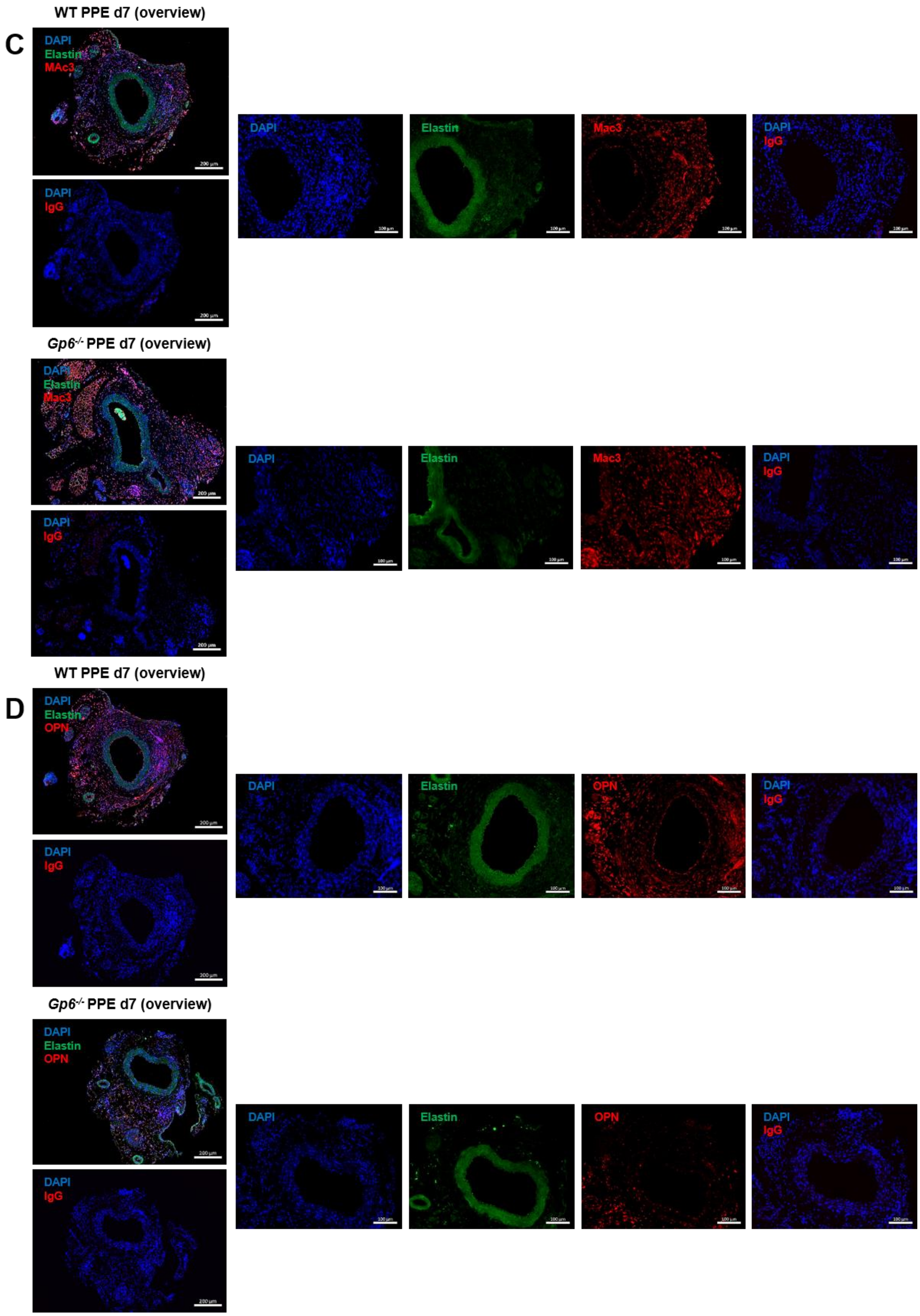
WT PPE d7 (overview)
C
DAPI
Elastin
MAc3
DAPI
IgG
DAPI
Elastin
Mac3
DAPI
IgG
Gp6-/- PPE d7 (overview)
DAPI
Elastin
Mac3
DAPI
IgG
DAPI
Elastin
Mac3
DAPI
IgG
WT PPE d7 (overview)
D
DAPI
Elastin
OPN
DAPI
IgG
DAPI
Elastin
OPN
DAPI
IgG
Gp6-/- PPE d7 (overview)
DAPI
Elastin
OPN
DAPI
IgG
DAPI
Elastin
OPN
DAPI
IgG

E

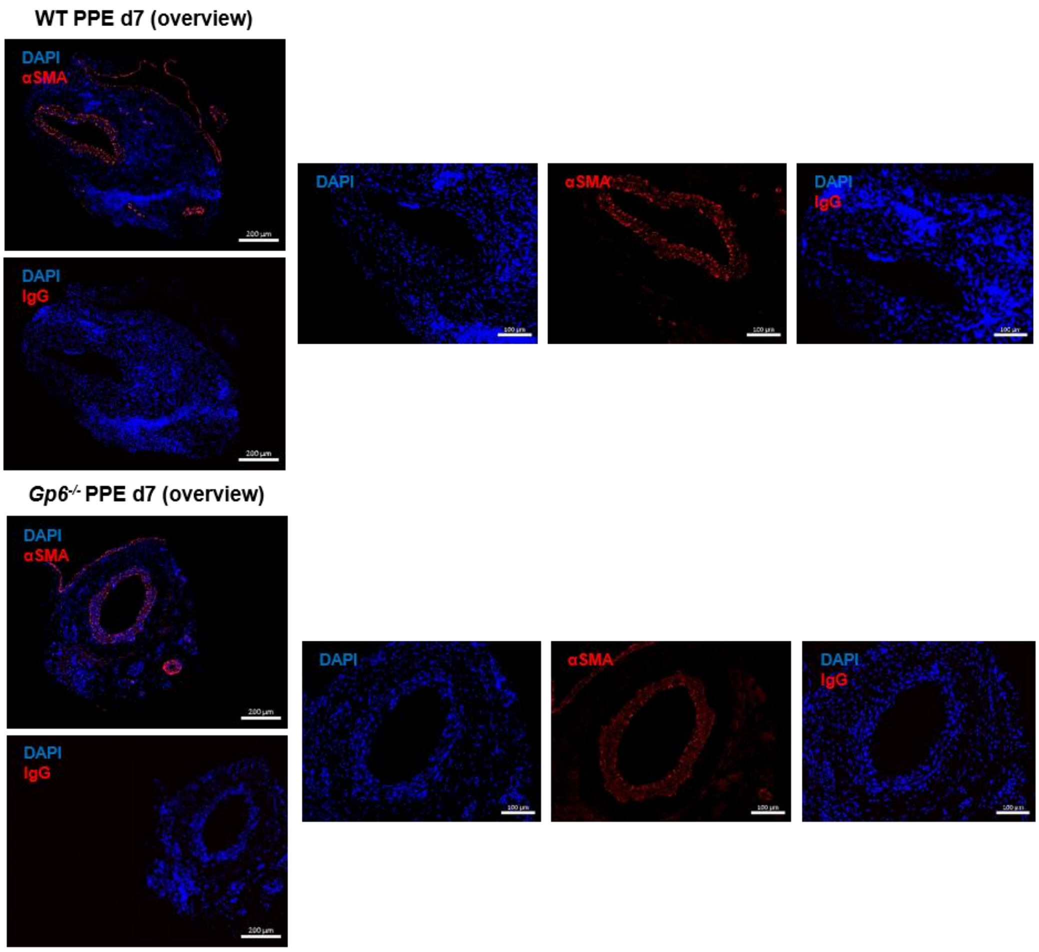
WT PPE d7 (overview)
DAPI
αSMA
DAPI
IgG
DAPI
αSMA
DAPI
IgG
Gp6-/- PPE d7 (overview)
DAPI
αSMA
DAPI
IgG
DAPI
αSMA
DAPI
IgG

F

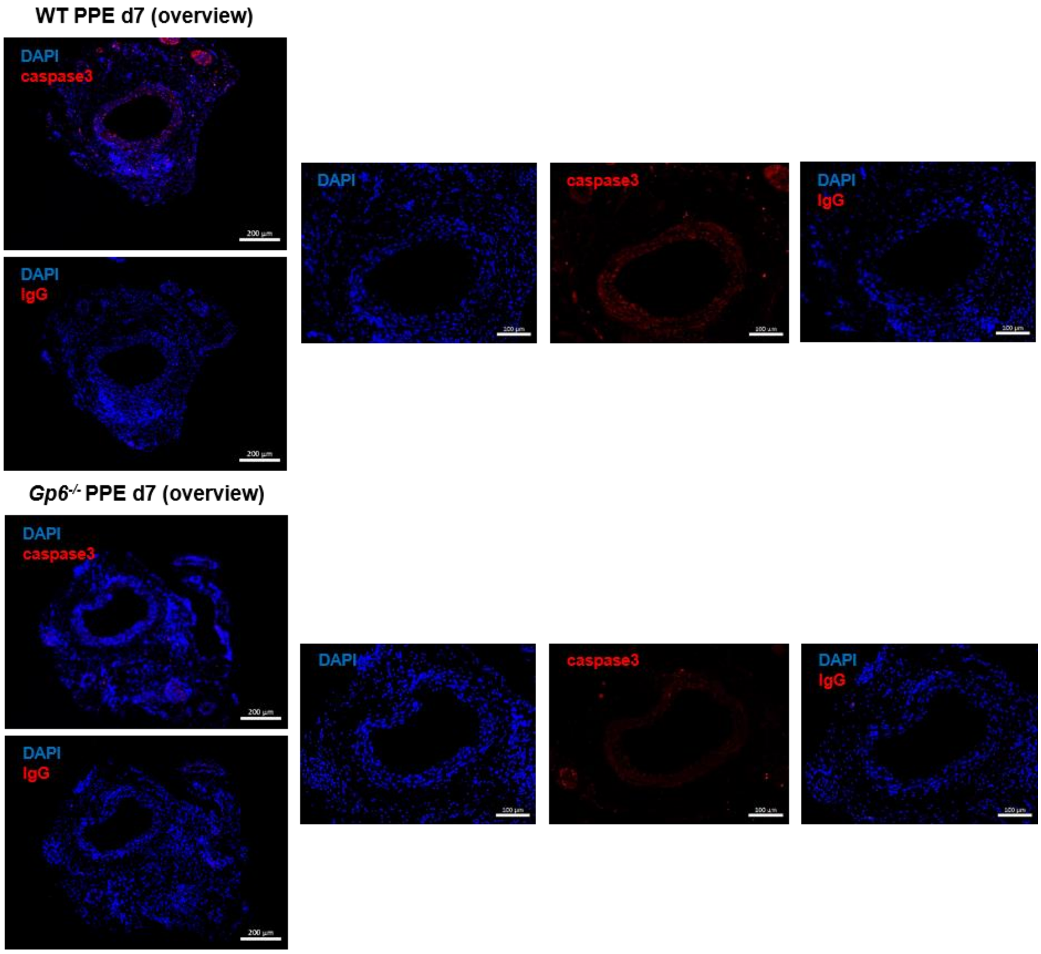
WT PPE d7 (overview)
DAPI
caspase3
DAPI
IgG
DAPI
caspase3
DAPI
IgG
Gp6-/- PPE d7 (overview)
DAPI
caspase3
DAPI
IgG
DAPI
caspase3
DAPI
IgG

**Figure S7. Immunofluorescence images of murine aortic tissue 7 days after experimental AAA induction.** (**A–F**) Representative immunofluorescence overview (left) and single-channel images (right) with respective IgG controls of murine aortic tissue from PPE operated WT (n=5–6) and *Gp6*$^{-/-}$ (n=4–5) mice at day 7. Aortic tissue was specifically stained for (**A**) neutrophils (anti-Ly6G/Cy3; red), (**B**) platelets (anti-GPIbα/Cy5; red), (**C**) macrophages (anti-Mac3/Cy5; red), (**D**) OPN (anti-OPN/Cy3; red), (**E**) α-SMA (anti- α-SMA/Cy3; red) or (**F**) active caspase3 (anti-cleaved caspase3/Cy3; red). Elastin autofluorescence (green) is shown and nuclei were stained with DAPI (blue). Scale bars: 200 µm (overview) and 100 µm. OPN = Osteopontin; PPE = Porcine pancreatic elastase (infusion); SMA = Smooth muscle actin.

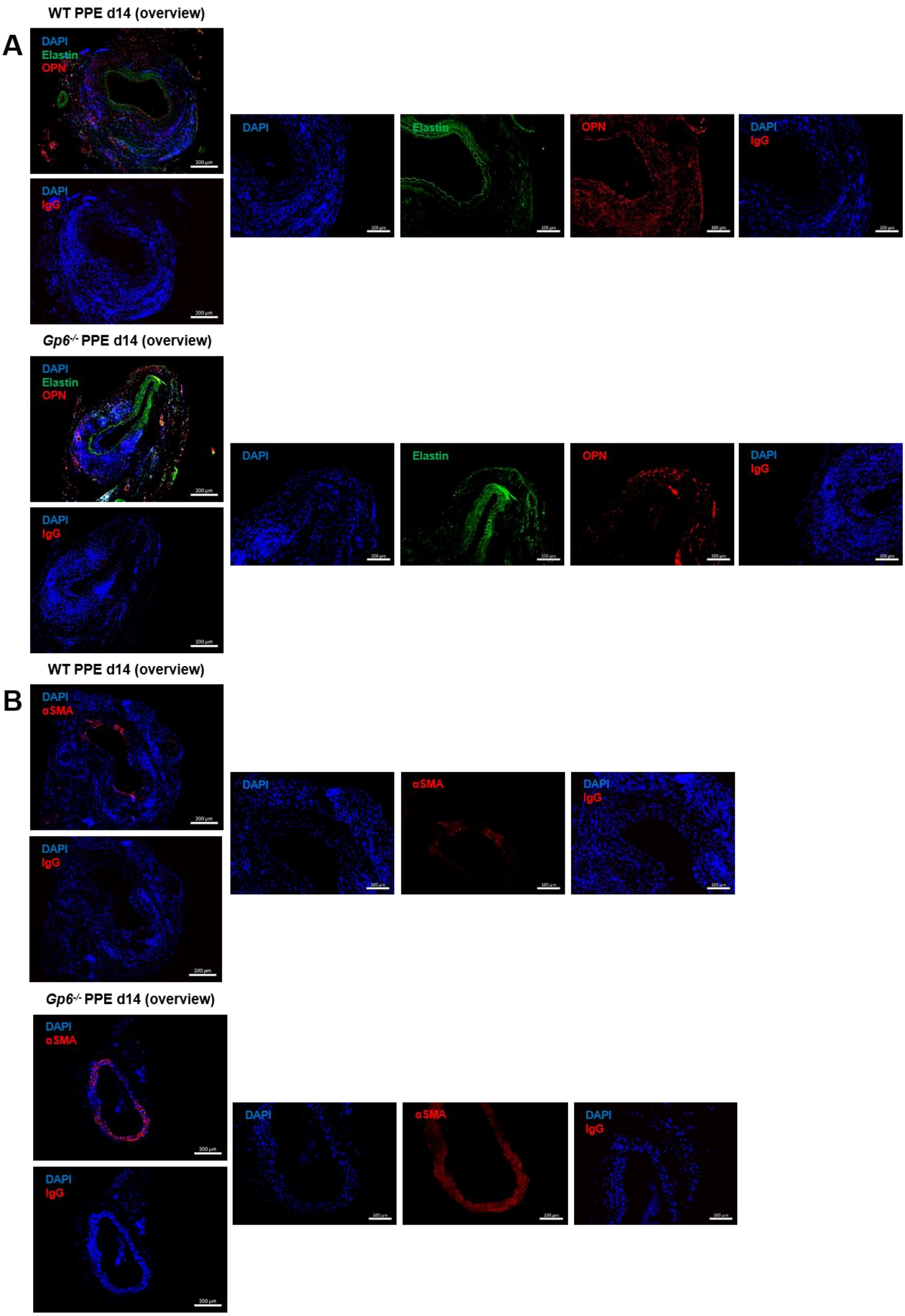
WT PPE d14 (overview)
A
DAPI
Elastin
OPN
DAPI
IgG
DAPI
Elastin
OPN
DAPI
IgG
Gp6-/- PPE d14 (overview)
DAPI
Elastin
OPN
DAPI
IgG
DAPI
Elastin
OPN
DAPI
IgG
WT PPE d14 (overview)
B
DAPI
αSMA
DAPI
IgG
DAPI
αSMA
DAPI
IgG
Gp6-/- PPE d14 (overview)
DAPI
αSMA
DAPI
IgG
DAPI
αSMA
DAPI
IgG

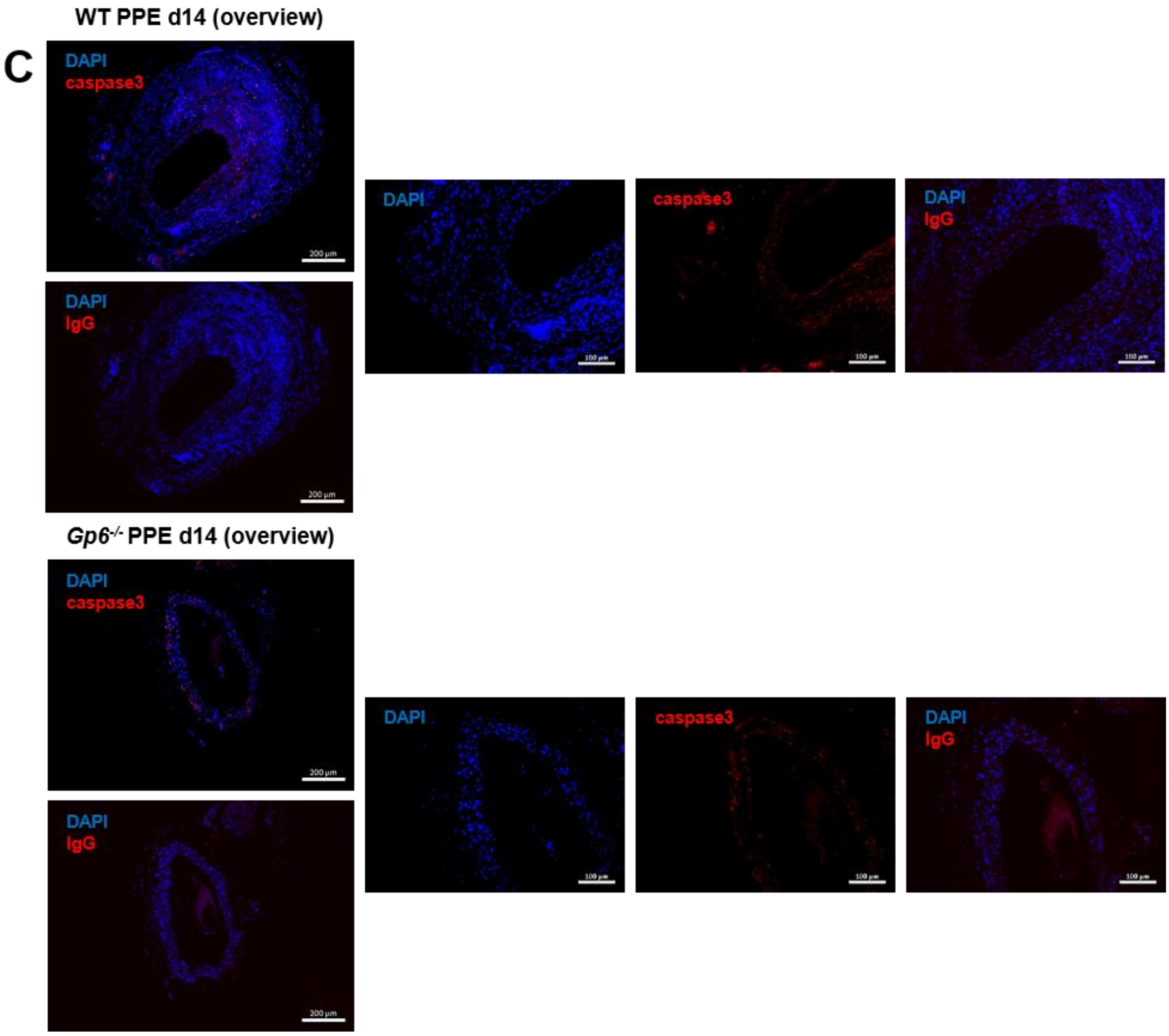


**Figure S8. Immunofluorescence images of murine aortic tissue 14 days after experimental AAA induction.** (**A–C**) Representative immunofluorescence overview (left) and single-channel images (right) with respective IgG controls of murine aortic tissue obtained from PPE operated WT (n=5–6) and *Gp6*$^{-/-}$ (n=4–5) mice at day 14. Aortic tissue was specifically stained for (**A**) OPN (anti-OPN/Cy3; red), (**B**) α-SMA (anti- α-SMA /Cy3; red) or (**C**) active caspase3 (anti-cleaved caspase3/Cy3; red). Elastin autofluorescence (green) is shown and nuclei were stained with DAPI (blue). Scale bars: 200 µm (overview) and 100 µm. OPN = Osteopontin; PPE = Porcine pancreatic elastase (infusion); SMA = Smooth muscle actin.

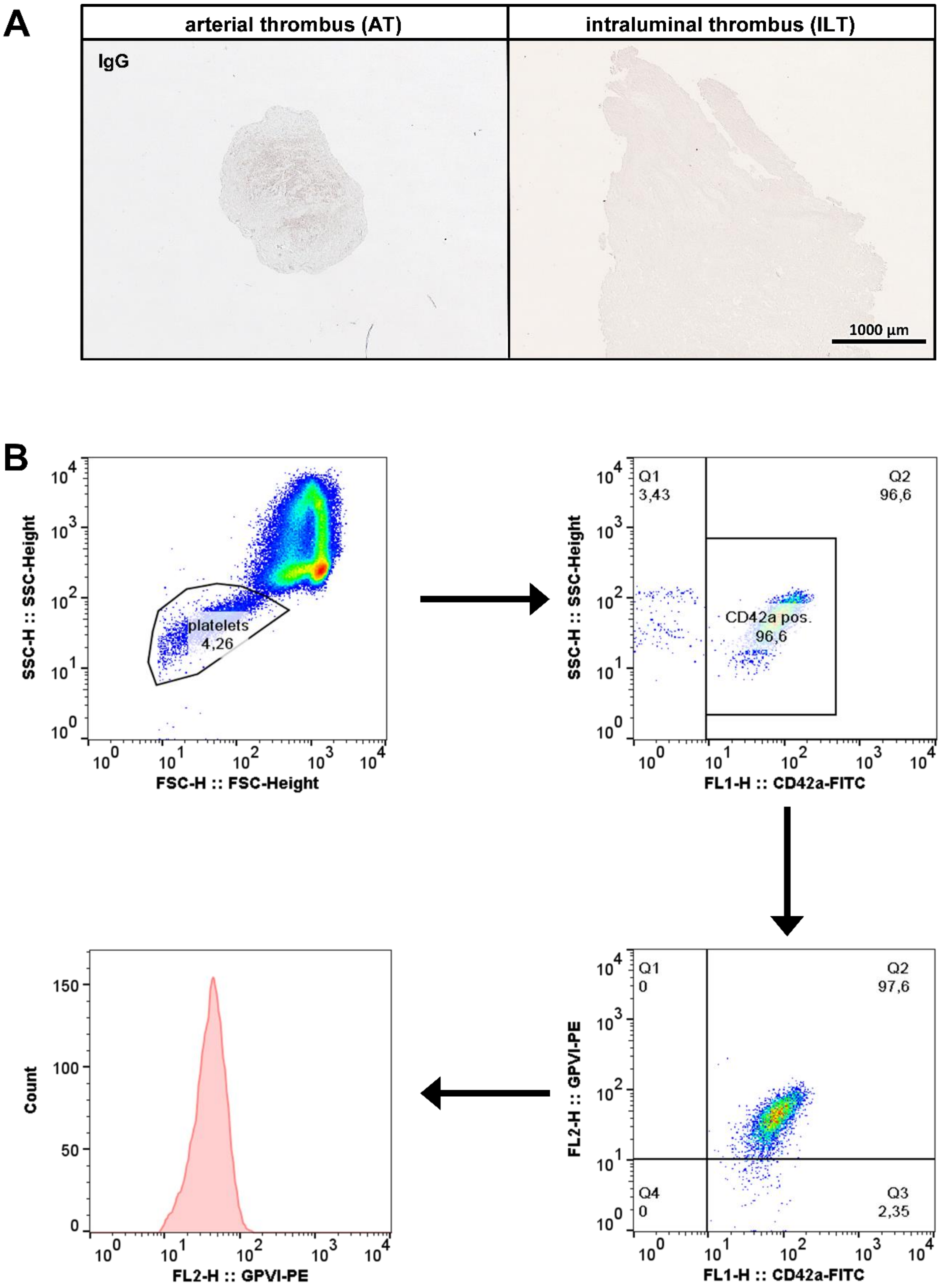


**Figure S9. Gating strategy for flow cytometric analysis of AAA patients.** (**A**) Representative images of IgG controls of immunohistochemically stained (platelets, anti-GPIbα) arterial thrombus (AT) tissue from thrombectomy patients (n=8), respectively intraluminal thrombus (ILT) tissue samples from AAA patients (n=12) samples. Scale bar: 1000 µm. (**B**) Gating strategy for analysing GPVI expression on platelets of AAA patients.

Whole blood samples were first gated on platelets according to their specific FSC/SSC profile. In addition, the platelet population was gated using the platelet specific marker GPIbα (CD42a) and a representative histogram for the GPVI expression on the platelet surface after gating is provided.
